# Mining Yeast Diversity Unveils Novel Targets for Improved Heterologous Laccase Production in *Saccharomyces cerevisiae*

**DOI:** 10.1101/2024.08.26.609787

**Authors:** Ryan Wei Kwan Wong, Marissa Foo, Jasmine R. S. Lay, Tiffany L. T. Wai, Jackson Moore, Fabien Dutreux, Cristen Molzahn, Corey Nislow, Vivien Measday, Joseph Schacherer, Thibault Mayor

## Abstract

The budding yeast *Saccharomyces cerevisiae* is a widely utilized host cell for recombinant protein production due to its well studied and annotated genome, its ability to secrete large and post-translationally modified proteins, fast growth and cost-effective culturing. However, recombinant protein yields from *S. cerevisiae* often fall behind that of other host systems. To address this, we developed a high throughput screen of wild, industrial and laboratory *S. cerevisiae* isolates to identify strains with a natural propensity for greater recombinant protein production, specifically focussing on laccase multicopper oxidases from the fungi *Trametes trogii* and *Myceliophthora thermophila*. Using this method, we identified 20 non-laboratory strains with higher capacity to produce active laccase. Interestingly, lower levels of laccase mRNA were measured in most cases, indicating that the drivers of elevated protein production capacity lie beyond the regulation of recombinant gene expression. We characterized the identified strains using complementary genomic and proteomic approaches to reveal several potential pathways driving the improved expression phenotype. Gene ontology analysis suggests broad changes in cellular metabolism, specifically in genes/proteins involved in carbohydrate catabolism, thiamine biosynthesis, transmembrane transport and vacuolar degradation. Targeted deletions of the hexose transporter *HXT11* and the Coat protein complex II interacting paralogs *PRM8* and *9*, involved in ER to Golgi transport, resulted in significantly improved laccase production from the S288C laboratory strain. Whereas the deletion of the Hsp110 *SSE1* gene, guided by our proteomic analysis, also led to higher laccase activity, we did not observe major changes of the protein homeostasis network within the strains with higher laccase activity. This study opens new avenues to leverage the vast diversity of *Saccharomyces cerevisiae* for recombinant protein production, as well as offers new strategies and insights to enhance recombinant protein yields of current strains.

## Introduction

*Saccharomyces cerevisiae* is a commonly used host cell for recombinant protein production in laboratory and industrial settings. While the comparatively low yields of *S. cerevisiae* are not often a barrier in laboratory settings, it has limited the use of this yeast species in large-scale industrial protein production applications. Still, *S. cerevisiae* has found applications in commercial production of several biopharmaceuticals such as insulin, growth hormones, factor XIII A-subunit (a blood clotting factor subunit), urate oxidase (treatment of high serum uric acid) and vaccines for hepatitis B and HPV [1]. Further increasing yields from *S. cerevisiae* would allow for its wider adoption as a recombinant protein host.

In our study we utilize the laccase multicopper oxidases, which belong to an enzyme superfamily with conserved plastocyanin-like domains and are used in diverse biotechnology applications. Laccases catalyze single electron oxidation of various phenolic and aromatic compounds with low substrate specificity [2]. The primary functions of laccases vary depending on the organism: some plants, like the namesake lacquer tree, use laccase for wound sealing [3]; insects, for cuticle hardening [4, 5]; bacteria, for melanization [6, 7] and the breakdown of lignin [8], the latter of which is one of the primary functions of laccases in fungi [9–12]. The promiscuity of these laccases for various aromatic compounds and their consideration as a “green” enzyme (i.e. one that produces water as the main by-product) explains why they have been extensively studied for their use in applications such as bioremediation, pulp and paper processing, wastewater treatment, food processing, biosensors, biofuel production and biofuel cells [13]. Despite their potential, these enzymes have often been difficult to express at high level [14].

Efforts have been made to improve recombinant protein yields from *S. cerevisiae* is an area of active research. The process of recombinant protein production has several potential bottlenecks at each stage of protein synthesis from transcription, translation and post-translational modifications, all the way through to secretion out of the cell. Attempts to improve the process are typically approached on a case-by-case, protein-by-protein basis. There are, however, a few common biological engineering targets, that can improve yields in a more general manner. For example, the endoplasmic reticulum (ER) resident proteins PDI and BiP are commonly overexpressed to increase protein secretion [15–17], while the vacuolar protease *PEP4* gene is commonly deleted to reduce aberrant degradation of the recombinant protein [18, 19]. Identification of new biological engineering targets have been accomplished by high-throughput screening of mutants [20], systematic screens of single gene deletions [21] and RNA interference libraries [22]. These large-scale screens have been largely restricted to a single strain background. Thus far, no large-scale screens of natural and industrial isolates have been performed, overlooking the vast diversity of *S. cerevisiae* strains and the growing annotation of the *Saccharomyces* pan-genome for recombinant protein production goals.

To expand the pool of strains used for recombinant protein production and identify new potential engineering targets, we used a library of ∼1000 strains (including laboratory, natural and industrial *Saccharomyces cerevisiae* isolates) to screen the ability of this diverse strain library to produce laccase in a high throughput expression assay. Specifically, we assessed the secretion of the fungal laccase enzymes originating from *Trametes trogii* (*ttLCC1*) as well as the *Myceliophthora thermophila* (*mtLCC1*) [23] to exploit the diversity of *S. cerevisiae* and its pan-genome. We first identified several strains with significantly improved laccase production ability compared to the laboratory strain BY4741 and then their genomes and proteomes. We then targeted several candidate genes that are absent or present at lower levels in the identified strains to find new pathways that could be manipulated to improve laccase production in the BY4741 lab strain. In this study we showcase the utility of leveraging the natural diversity of *Saccharomyces cerevisiae*, via high throughput functional screening of non-laboratory strains, for recombinant protein production.

## Results

### A Novel High Throughput Screen of Natural and Domesticated *Saccharomyces cerevisiae* Isolates

To exploit the vast diversity of *Saccharomyces cerevisiae* strains for recombinant protein production applications, we developed a novel screening pipeline using production of laccase from *Trametes trogii* (*ttLCC1*) as a test case. Because most of the strains tested lack auxotrophic markers, we generated an expression vehicle comprising of a *CEN6/ARSH4* plasmid with a dominant selectable *kanMX6* marker and *ttLCC1* under the control of the strong constitutive *GPD1* promoter and *CYC1* terminator (Fig. S1A). We confirmed the expression and secretion of laccase in the BY4741 reference strain by assaying its activity in the media after removal of the yeast cells using a colorimetric assay based on the substrate ABTS (2,2’-azino-bis(3-ethylbenzothiazoline-6-sulfonic acid) [24]. We observed a peak of activity after 4 days, soon after the culture reaches saturation in our growth conditions (Fig. S1B). Next, we tested if laccase activity could be assessed in high-throughput with cells grown in 96-deep-well plates, confirming that well position does not skew laccase activity (Fig. S1C). With screening conditions optimized in a 96-well format, we proceeded with transformation and screening of the strain library.

We assessed the 1000 strain library that was previously sequenced [25], supplemented with additional *S. cerevisiae* isolates from the Okanagan wine region and from our own collection. Because 73 strains already contain the kanamycin resistance marker, we could only assess a total of 981 strains. Following transformation with our *CEN* plasmid, we obtained 597 transformed strains for expression of the ttLcc1 laccase (Additional file 1). Transformation of non-laboratory strain may be impacted by several genes and cell wall structure [26], likely explaining why only ∼60% of the strains were successfully transformed. The ecological origins of the transformed strains display a good representation of the whole library (Fig. S1D & E). The transformed strains were arrayed into 96-well plates and single replicates of each strain were used for the initial screening by inoculating each cell population at the same density. An internal control consisting of a BY4741 replicate, was included in each plate. After 4 days, the media, after removal of all cells, was assessed by the ABTS assay to determine laccase activity (Fig. 1A). From this initial screening, 47 strains were identified as preliminary hits using a threshold of laccase activity of at least 3 median absolute deviations (MAD) above the median activity of all strains screened (Fig. 1B; Additional file 1). To confirm that these strains produce the recombinant proteins at a higher level, we re-transformed them with the *ttLCC1* laccase plasmid and re-arrayed them into 96-well plates with 4 biological replicates per strain. In this second screening, 9 of the strains were identified to have significantly greater ttLcc1 activity than BY4741 (Fig. 1C). These strains constituted a first set of isolates to be further characterized.

**Fig. 1.**
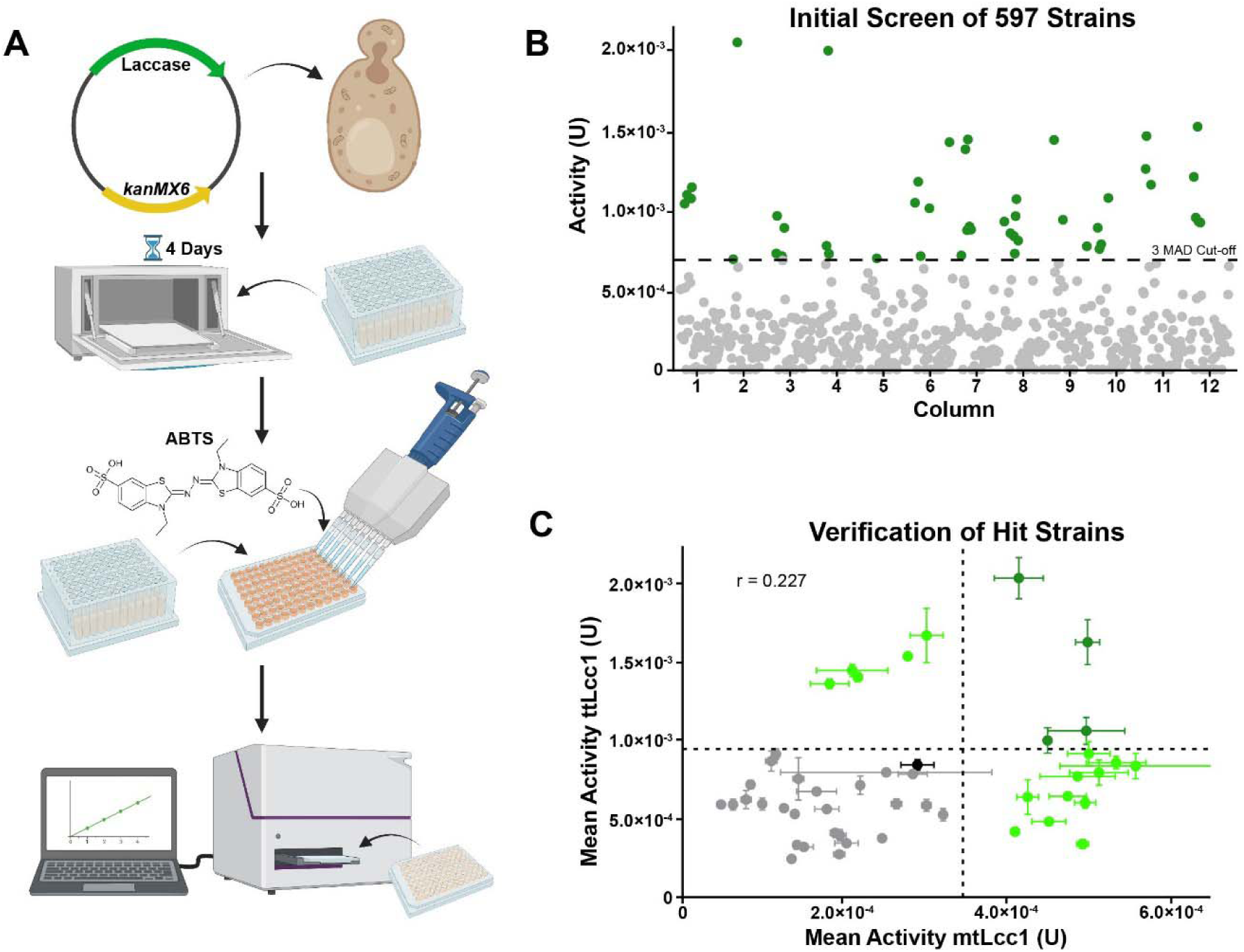
**A)** Workflow of the high-throughput laccase production screen. **B)** Plots of ttLcc1 laccase activity for ∼600 assessed strains (enzymatic activity Units). 47 hits above the 3 M.A.D. cut-off are shown in green. **C)** Secondary screening with *ttLCC1* (x-axis) and *mtLCC1* (y-axis; mean enzymatic activity Units) using the 47 preliminary hits (n=4). Validated hits are shown in green, reference (BY4741) in black and significance cut-offs as dashed lines. Pearson correlation of the two laccase activities in assessed strains is shown.

To determine if ttLcc1 expression was extensible to other recombinant proteins, we tested a laccase that was identified in *Myceliophthora thermophila* (mtLcc1). While both ttLcc1 and mtLcc1 are laccases with 3 plastocyanin-like copper binding domains, they share only 27.6% identity and 39.2% similarity at the protein sequence level (Fig. S1E). The activity of mtLcc1 was assessed in parallel with ttLcc1 in the 47 strains isolated in the first screen. Surprisingly, there was a relatively low correlation of the measured activities of the two laccases between the strains, with a Pearson correlation co-efficient of 0.227 (Fig. 1C). Nonetheless, 15 strains had significantly greater mtLcc1 activity than BY4741, including 4 strains with significantly greater activity for both laccases, resulting in a total of 20 strains producing higher laccase activity. These results indicate that the ability for a strain to overexpress one laccase does not necessarily correlate to higher production of another related enzyme. Nevertheless, whatever biological features these strains have to allow them to produce either protein better than BY4741 can point to new targets that may be leveraged by additional strain engineering.

### Laccase Production Affinity is Present in a Diversity of Strain Origins

A major differentiation of these strains compared to BY4741, is the environment or application that these strains have adapted to. Thus, we next looked at the origins of the hit strains to determine their relatedness and better understand how their evolutionary adaptations might potentially affect heterologous protein production. To assess the relationships between the 20 strains identified as hits for ttLcc1 and mtLcc1, we constructed a phylogenetic tree using a previously published distance matrix [25], annotating the origins of each clade and highlighting our hit strains. The hits are distributed in 5 different clades encompassing the Wine/European (8 strains), French dairy (3 strains), African beer (2 strains), Mixed origin (6 strains) and Mosaic region 3 (1 strain) clades (Fig. 2). W303, a strain that closely clusters with S288C (the parent of BY4741 [27]) due to sharing 85.4% genomic identity and a shared ancestor [28], is also found in the Mosaic region 3 clade and was used as the reference strain because BY4741 is not present in the sequencing dataset of these strains [25]. There are 4 pairs of strains found on neighbouring branches (namely A-14 and D-1, DBVPG1374 and DBVPG1714, CLIB650 and CLIB655, and CCY_21-4-97 and CCY_21-4-98), indicating that the traits that allowed for higher laccase production may have originated in recent common ancestor strains. In agreement with this hypothesis, the closely related CLIB650 and CLIB655 strains are strong producers of ttLcc1, whereas DBVPG1374 and DBPVG1714 are strong mtLcc1 producers. Overall, there is considerable genetic distance between strains that exhibited improved laccase production, indicating high diversity among the identified strains.

**Fig. 2.**
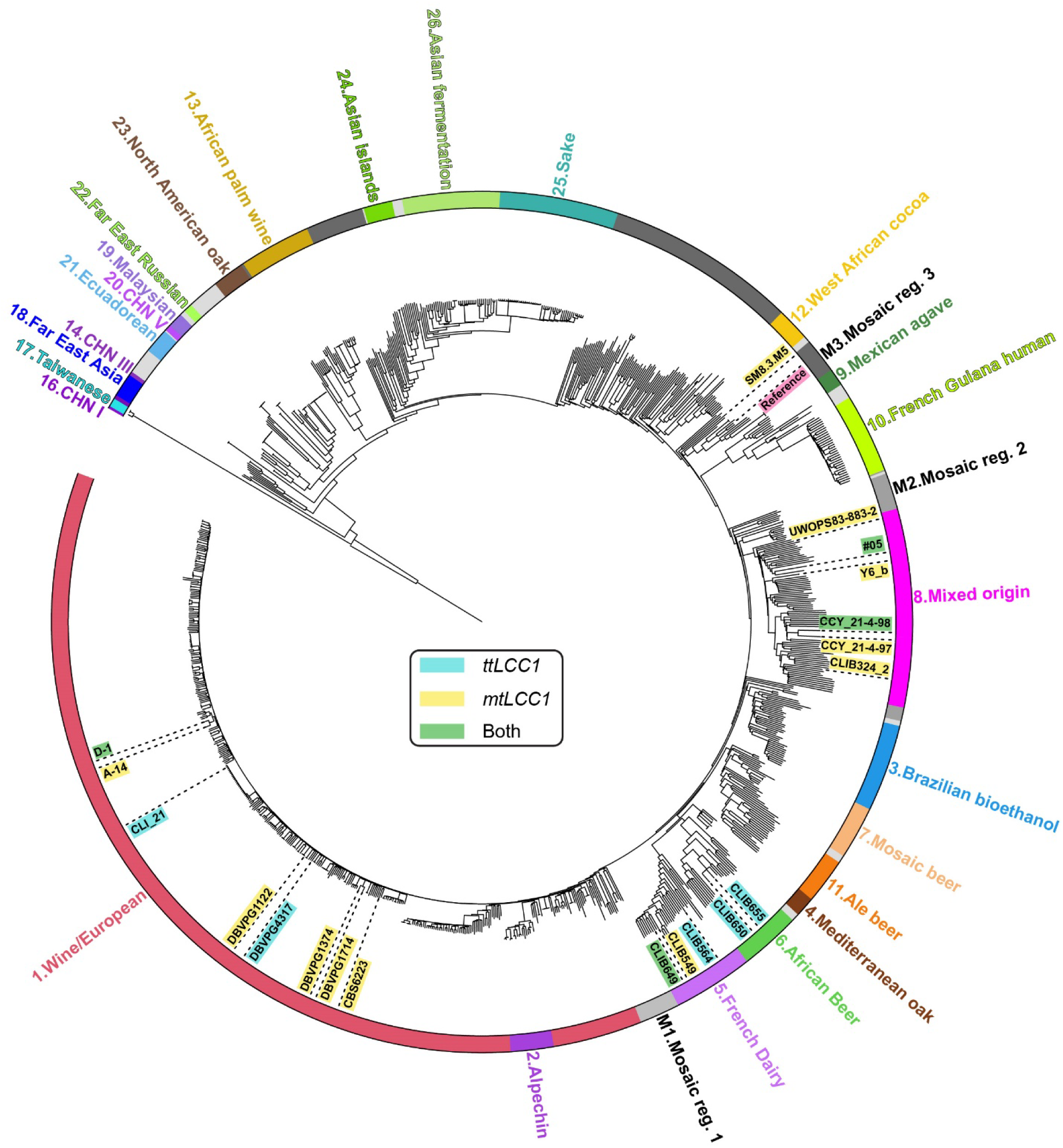
Phylogenetic tree highlighting the hit strains and the phylogenetic clade to which they belong. Strains are highlighted according to the laccase for which they were a hit. The W303 reference strain that closely clusters with BY4741 is also highlighted.

### Determining the Level of Organization of Laccase Production Improvement

With such strain diversity, it is possible that the mechanisms controlling the increase in laccase production may be diverse. For instance, greater growth may lead to increased laccase production by increasing the biomass available for production. While there was variation in optical densities (OD_600_) that relate to the cell biomass, ranging from -5% to +24% and that may be impacted by cell growth, the variation in laccase activity was greater, ranging from +19% to +143% (Fig. S2A). This suggests that increased biomass is not a major factor in the observed increase in laccase activity. Timing of laccase production did not appear to be a major factor, although it may slightly vary depending on the strain (Fig. S2B). For instance, the activity of the laccase reached its maximum activity in CLIB649 after 3 days. At the nucleic acid level, transcription is a limiting factor of protein production with more mRNA transcripts allowing more translation of a given protein. To determine if transcription is a ttLcc1 expression enhancing factor for our strains, we conducted reverse transcriptase-quantitative PCR (RT-qPCR) to assess the relative amount of *ttLCC1* mRNA and compared them to laccase activity from the same strains. *UBC6* was used as the internal reference as its expression had been demonstrated to be stable in different conditions and strains (Fig. S2C) [29]. Specificity of the RT-qPCR was assessed by melting curve to ensure a single target was amplified (Fig. S2D). Except for two strains with higher *ttLCC1* mRNA levels (CLIB564 and D-1), one strain (DBVPG4317) had no significant difference in *ttLCC1* mRNA levels, and the remaining six strains had significantly lower *ttLCC1* mRNA levels compared to the BY4741 reference strain (Fig 3A). We next compared the laccase activity measured in parallel in these same strains and normalized to the *ttLCC1* mRNA levels. In all but two strains (CLIB564 and D-1), the normalized laccase activity was significantly higher relative to the reference strain (Fig. 3B). These findings suggest that the elevated laccase activity in the assessed strains is likely due to changes affecting protein homeostasis and not plasmid copy number or elevated transcription.

**Fig. 3.**
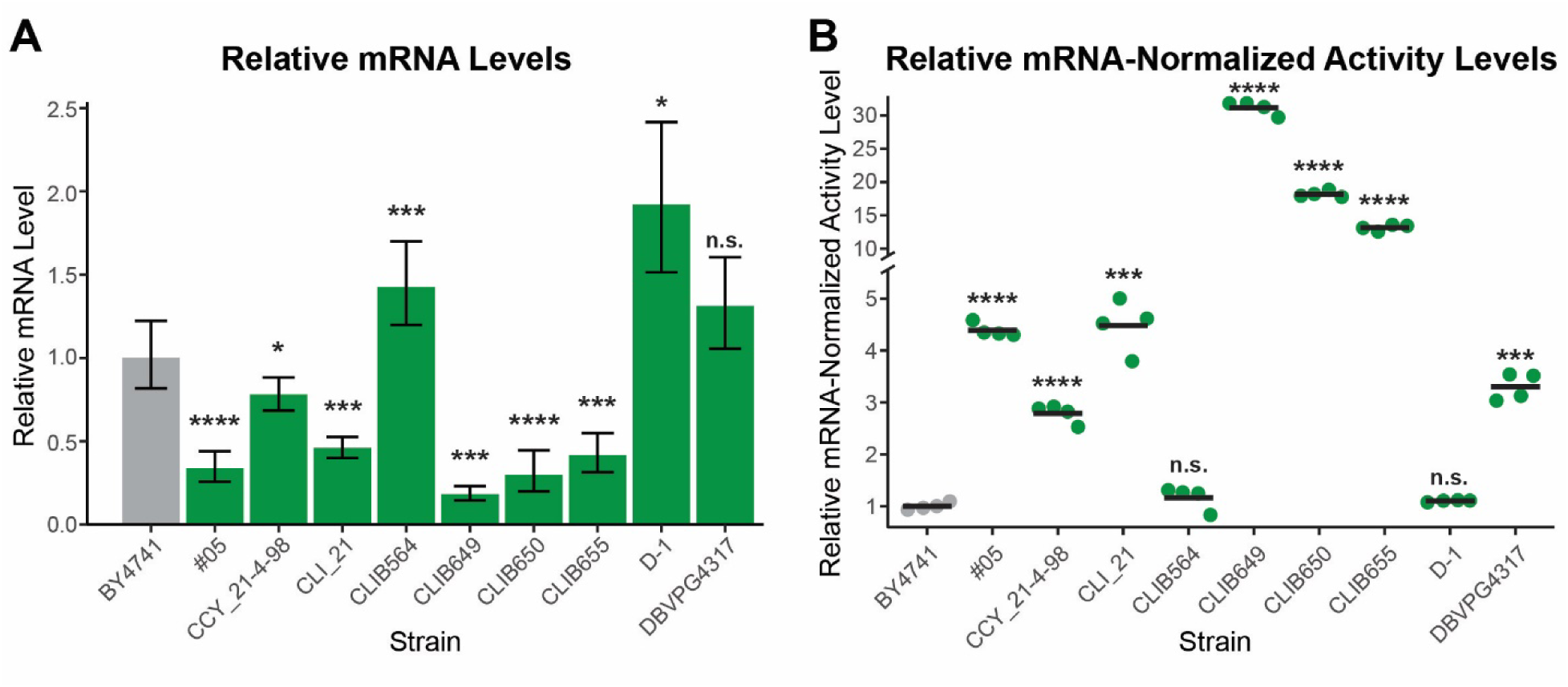
**A)** Relative *ttLCC1* mRNA levels of the indicated strains. After 4-days growth, an equivalent of 25 OD_600_ of cells were processed for mRNA purification and RT-qPCR (n=3). Levels of *ttLCC1* mRNA were normalized to *UBC6.* **B)** Laccase activity relative to the averaged *ttLCC1* mRNA levels for each strain in shown. Laccase activity was measured in parallel to mRNA quantitation (p-values: * < 0.05, ** < 0.01 *** < 0.001, **** < 0.0001).

### Characterization of Hits using Genomics

To better understand what may contribute to the improvements in laccase production, we conducted genome wide association studies (GWAS) and open reading frame (ORF) enrichment and depletion analyses utilizing published sequencing data [25]. GWAS was conducted using the laccase activity phenotypes for both single nucleotide polymorphisms (SNPs) and copy number variations (CNVs) using activities from all transformed strains. A single significant SNP was found in *ATG27* (a membrane protein involved in autophagy, also known as *ETF1*) (Fig. S3A) and a single significant CNV was found encompassing YPL273W and YPL274W, *SAM4* (an S-adenosylmethionine-homocysteine methyltransferase) and *SAM3* (an S-adenosylmethionine permease) respectively (Fig S3B). With meagre hits from GWAS, we decided to identify ORFs that were either enriched or depleted within the hit strains in comparison to the remaining library using a Fisher’s exact test. This analysis, which included the *S. cerevisiae* pan-genome, identified 23 ORFs as enriched among the 20 identified strains (p-value <0.05), 10 ORFs were significantly depleted but had a compensatory enrichment of a highly similar ORF, and 38 ORFs were depleted (Fig. 4A). The highly similar ORFs, which are indicated as ‘ORF-like’ in Fig 4A, are genes that are not present in the S288C reference genome but have been identified by sequencing of the diverse *S. cerevisiae* strains and are present in the *S. cerevisiae* pan-genome [25].

**Fig. 4.**
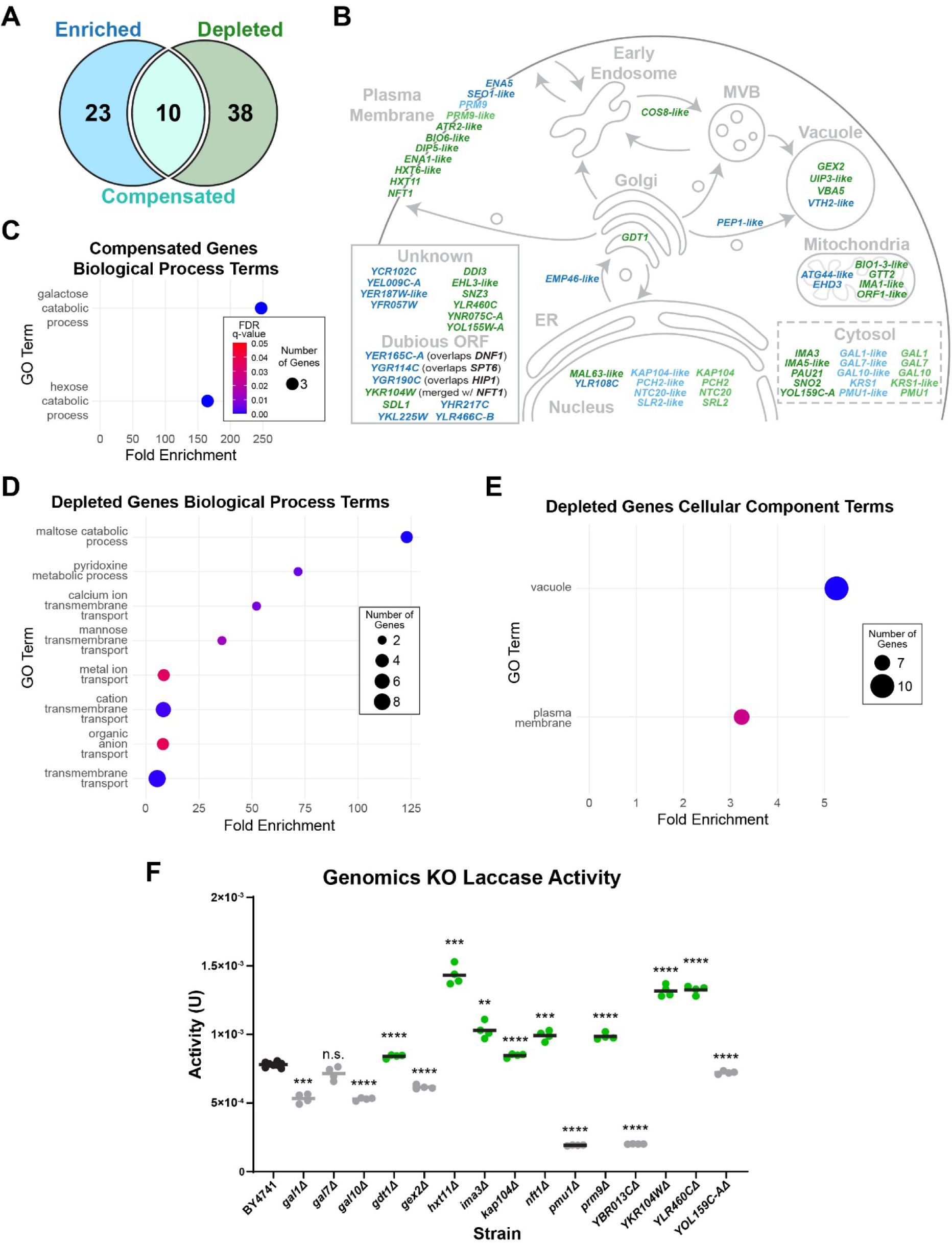
**A)** Number of ORFs that are either significantly enriched or depleted in the 20 strains with higher laccase activity. We first identified genes that are either present or absent in any of the 20 hit strains in comparison to the reference strain BY4741, and then used Fisher’s exact test to determine whether these genes are statistically enriched or depleted in comparison to the whole strain library. The compensated group consists of 10 depleted genes for which a similar ORF is also enriched. **B)** Cell map of the localization of known genes or of their close homolog. Enriched ORFs shown in blue, depleted in green and compensated lighter colours. **C, D and E)** Dot plots of the compensated and depleted genes groups, indicating Biological Process or Cellular Component GO terms enriched in each group. The scale of the false discovery rate (FDR) q-values are shown in C. **F)** Select gene targets from genomic analysis were knocked out and tested for their effect on ttLcc1 activity after 4-days growth. BY4741 was used as the control and reference for statistical tests. Significantly higher or lower laccase activities indicated in green or in grey, respectively (p-values: * < 0.05, ** < 0.01 *** < 0.001, **** < 0.0001).

To obtain an overview of which pathways may be involved, we first mapped the enriched and depleted ORFs according to the cell localization of the corresponding proteins or related proteins (Fig. 4B). Many depleted genes encode for proteins that are localized in the plasma membrane, as well as in the vacuole. Absence of these genes may impact the secretion of their recombinant protein. Interestingly, nuclear and cytosolic localized ORFs depleted in the identified strains are often compensated with a similar ORF enrichment. We assumed that this is a faithful compensation with the enriched gene able to fulfill the functions of the depleted. Several ORFs do not have any assigned cell localization or are annotated as dubious ORFs. Notably, dubious ORFs are more likely to be enriched rather than depleted.

We next performed Gene Ontology (GO) term enrichment analysis on the ORFs significantly enriched or depleted among the identified strains. To facilitate the analysis, the ORF sets were manually curated by replacing gene-like annotations to only include ORFs of characterized genes. From the group of enriched genes, we found no significant GO term enrichment. This was expected as many of the enriched ORFs are dubious and not expected to encode a functional protein. From the compensated gene group, the GO terms of biological processes linked to galactose and hexose catabolism are enriched (Fig 4C), driven by *GAL* genes that are encoded by homologous genes in many strains. From the depleted genes group, biological process terms covering hexose metabolism and transmembrane transport are enriched (Fig. 4D). Several *IMA* and *HXT* genes, involved in hexose catabolism and transport, are significantly depleted among the identified strains, as well as *SNZ3* and *SNO2* that are involved in pyridoxine (vitamin B6) and thiamine biosynthesis. Depletion of these genes suggest that rewiring of metabolic pathways could impact heterologous production [30]. In agreement with the depletion of genes linked to transmembrane transport, plasma membrane localization is also significantly enriched among this group of depleted genes (Fig 4E). In addition, vacuole localization is enriched with 10 genes mapping to that term. One possibility is that depletion of vacuole genes could play a role in diverting proteins from vacuole degradation leading to higher secreted laccase activity.

### Validation of Genomic Hits

With numerous genes depleted within the identified strains, we sought to validate a subset of these ORFs by knocking out (KO) these genes in our reference strain BY4741. We generated 15 KOs by replacing each gene with a *HIS3* cassette and assayed ttLcc1 activity in each mutant, including six genes for which a corresponding homologous ORF is enriched (i.e., “compensated” group). Eight out of 15 deletions resulted in significantly higher laccase activity compared to BY4741 (Fig. 4F). We verified that the insertion of *HIS3* itself (in this case into the *HO* locus) did not cause laccase activity to increase. It in fact, mildly decreased activity when tested on 11 unique isolates (Fig. S4). Of these KOs, *gdt1*Δ had the least improvement to laccase activity, increasing activity by a modest 8% (Fig. 4F). We initially postulated that in each case of compensation, the enriched gene effectively would replace the depleted homologous gene. Thus, these genes were not expected to give significantly increased laccase activity. Contrary to our expectations, KOs targeting *KAP104* and *PRM9*, two of the “compensated” genes, resulted in significantly higher laccase activity, close to 10% and 25%, respectively. The best performing KOs were *hxt11*Δ, *ykr104w*Δ and *ylr460c*Δ. Deletion of these targets had the greatest improvement in laccase activity, nearly 85%, 70% and 70%, respectively (Fig. 4F). *HXT11* is a broad-spectrum hexose transporter capable of transporting glucose, fructose, mannose and galactose. *YLR460C* is an uncharacterized member of the quinone oxidoreductase family. In S288C, *YKR104W* is an ORF generated due to a stop codon of the adjacent ORF, *NFT1/YKR103W*, resulting in the 3’ truncation of *NFT1. YKR104W* encodes for an ATPase domain of the full length *NFT1* gene that is a merge of *YKR103W* and *YKR104W* in other strain backgrounds. Our results indicate that several genes absent in the identified strains may, at least partially, explain why higher laccase activity is observed in these cells.

### Characterization of Hits *via* Proteomics

To complement our genomic analysis, we assessed the proteome of the 20 strains with higher laccase activity using data-independent acquisition mass spectrometry. Notably, the protein mass spectrometry analysis can provide further insights into differences between the identified strains that cannot be easily predicted using genomic information. Cells from 4 biological replicates were collected after 4 days of growth and laccase expression for an analysis of the whole cell lysates. Mass spectrometry analysis of additional technical replicates of one reference sample was performed throughout the experiment to verify the homogeneity of the analysis, and a total of 14 samples were analyzed (Additional file 2). On average, we quantified 3,708 proteins per strain for a total of 4,021 distinct quantified proteins. We verified that there is a high correlation of the quantified proteins between biological replicates, with only three samples with an R^2^ value below 0.85 when compared to all its other biological replicates that we removed from the analysis (Fig. S5A). To further evaluate the proteomic results, we compared the protein intensities within the BY4741 technical replicates (n=14), which display low coefficient of variances (CVs) with a median CV of ∼8%, confirming the good quality of the measurements (Fig. S5B). The median CV increases to 22% when considering, instead, BY4741 biological replicates (Fig. S5C; n=7). The increase of the CVs is more marked for proteins with lower signal intensities (Fig. S5D), indicating some stochastic events are more likely to happen for low abundant proteins (either at the expression or proteolysis level) during the growth of the cells for 4 days. Comparably, the median CVs between biological replicates of the other strains range from 11% to 28% indicating some changes in the variability between biological replicates depending on the strains (Fig. S5E). Importantly, the CVs are in most cases lower in comparison to the reference strain. Several characterized strains display strong correlation of the quantified values when compared between each other, indicating their proteomes are very similar (Fig 5A). However, in some cases, the R values are much lower (e.g., when comparing BY4741 and CLIB655). We examined more closely the averaged protein intensities of two representative strains (CLI_21 and CLIB655). Interestingly, the variability of the measured intensities is well spread out in comparison to the lab reference strain (Fig S5F), indicating the variability affects both low and high expressed proteins. This is surprising, especially since recent work shows that proteomes display a good buffering capacity between different isolated strains [31, 32]. To better visualize the relationship between the different strains, we performed a principal component analysis (PCA) of all the analyzed samples which showed 5 visible groups of strains (Fig 5B). The PCA data demonstrates that BY4741 and CLIB324_2 are clearly separated in their own groups, while CLIB650 and CLIB655 are grouped together (Fig 5B). These 3 groups are the most separated from the others indicating distinct proteomic signatures. The two other groups each comprise a larger number of strains that are not markedly different from each other. The neighbouring pairs of A-14 and D-1, DBPVG1374 and 1714, CLIB650 and 655, and CCY_21-4-97 and 98 in the phylogenetic tree are all found clustering closely with their partner. Likewise, the closely related CLIB549 and 649 also closely cluster. These results show that, perhaps not surprisingly, more closely related strains share a closer proteome signature. As well, the separation of the BY4741 proteome from the other strains is consistent with the S288C strain being a phenotypic outlier when compared with diverse *S. cerevisiae* strains for over 600 traits [33].

**Fig. 5.**
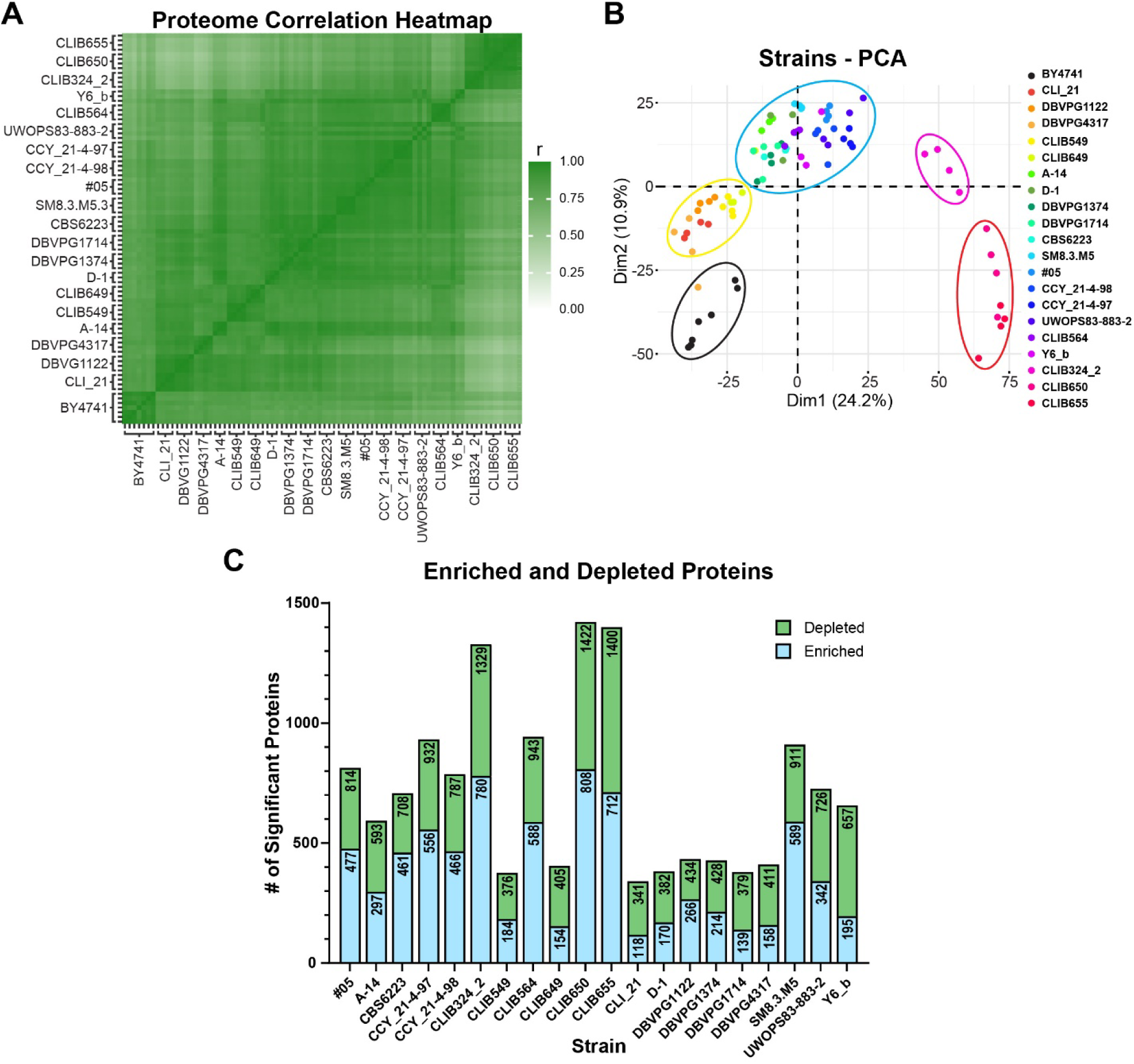
**A)** Heatmap representing the Pearson correlation of measured protein intensities of indicated strains compared to all other collected proteomes. **B)** Principal component analysis of the individual proteomes of the 20 hit strains and BY4741. The proteomes form 5 groups, highlighted by coloured ovals. **C)** Stacked bar chart of significantly enriched and depleted proteins from each strain after comparing with the reference BY4741 strain.

We next compared the abundance of individual proteins in the whole cell lysates of the 20 strains with the BY4741 lab strain. On average, 719 proteins displayed a significant two-fold change, ranging from 341 to 1,422 differentially expressed proteins in CLI_21 and CLIB650, respectively (Fig. 5C). In total, we detected 1565 differently enriched and 1481 depleted proteins across the 20 strains individually compared to BY4741 (Fig. S5G). We included in the search ttLcc1 that was expressed in these cells and observed significantly lower levels of the recombinant protein in the whole cell lysates of the 9 strains that produce higher activity of secreted ttLcc1 (Fig. S5H). This may be explained by a lower dwelling time of the recombinant protein during cell trafficking, resulting in more efficient secretion. Among the enriched proteins, seven are encoded by genes accordingly enriched in the genomics analysis (Atg44-like, Ehd3, Gal1-like, Gal7-like, Gal10-like, Krs1 and the EC1118_1O4_6502g uncharacterized and hypothetical amino acid transporter), whereas 10 depleted proteins are encoded by depleted genes in the genomics analysis (Ddi3, Gal1, Ima3, Ntc20, Pau21, Pmu1, Srl2, Snz3, YOL159C-A and YRL460C) (Fig. 4B).

To focus our efforts, we filtered proteins affected in at least 50% of the hit strains representing 239 enriched and 212 depleted proteins (Fig S5G). The GO analysis revealed that metabolic processes are enriched among proteins expressed at higher levels in the assessed hit strains, including “small molecule metabolic process”, “cellular amino acid metabolic process” and “ribonucleotide metabolic process” (Fig. 6A). Each of these three terms encompasses over 100 proteins, whilst the general term “metabolic process” encompasses 215 out of 239 enriched proteins, indicating that a main difference in the proteome of the hit strains compared with the reference strain is a distinct wiring of metabolic pathways. These proteins are also associated with cellular components terms related to mitochondria, proteasome, stress granule and nucleolus (Fig. 6B), possibly underscoring a higher capacity to respond to stresses such as heterologous protein production. Depleted proteins are significantly associated with terms related to catabolic processes (Fig. 6C). Interestingly, carbohydrate catabolic process related terms are also associated with depleted genes (albeit driven by different proteins/genes). Cellular component terms include vacuolar lumen, cell surface and extracellular region (Fig. 6D). The vacuolar lumen term is also associated with depleted proteins, as for depleted genes, suggesting a possible role in reduced vacuolar degradation (Fig. 4D and 6D).

**Fig. 6.**
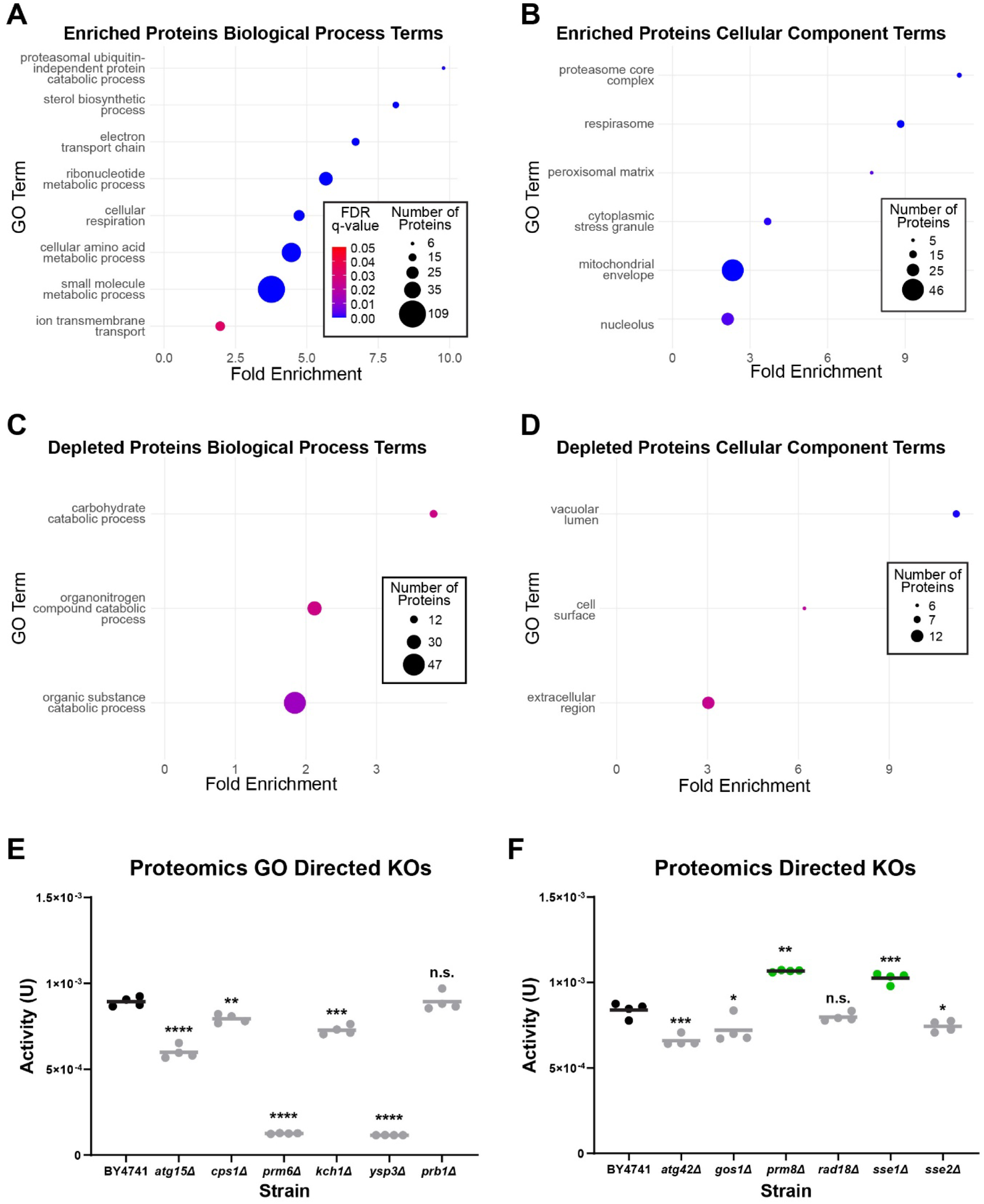
**A to D)** Dot plots of the Biological Process and Cellular Component GO terms enriched in the affected protein groups. FDR q-value scale as in A. **E and F)** Laccase activity after 4-days growth in the indicated KO strains selected following the proteomic analysis. Targets were selected based on GO term enrichment (E) or associated biological pathway of interest (F). BY4741 was used as the control and reference for statistical tests. Significantly higher or lower activities indicated in green or grey, respectively (p-values: * < 0.05, ** < 0.01 *** < 0.001, **** < 0.0001).

### Assessment of the Impact on Laccase Production of Affected Proteins

We next sought to select a few targets shown to be depleted in the proteomic analysis to assess their potential impact on heterologous protein production. We selected several targets based on GO term enrichment, namely, *ATG15*, *CPS1*, *PRM6* and *YSP3,* which are all found in the vacuole. In addition, we selected targets involve in vesicle trafficking, protein folding and homeostasis: *ATG42*, *GOS1*, *PRM8*, *RAD18* and *SSE2.* We also targeted for deletion *KCH1, PRB1* and *SSE1* that are paralogs of *PRM6, YSP3 and SSE2*, respectively. A total of 12 KOs were generated and tested for *ttLCC1* activity. The GO directed individual KOs did not result in any significant improvement to laccase activity (Fig. 6E). While not essential, these full knockouts may have a lower ability to scavenge nutrients during extended incubation time [34–37]. Nonetheless, we detected significantly higher laccase activity in *prm8*Δ (+27%) and *sse1*Δ (+22%) (Fig. 6F). *PRM8* is the paralog of *PRM9*, a hit we validated in the genomic analysis, and has roles in coat protein complex II (COPII) binding [38] in the early stages of secretion. We assessed both strains side by side to show that deletions of *PRM8* and *9* had similar impact on laccase activity (Fig. S6). *SSE1* has roles in protein folding being an Hsp110 with nucleotide exchange factor activity on the Ssa and Ssb family Hsp70s, as well as acting as a co-chaperone of the Hsp90 chaperone complex [39, 40]. These results indicate that changes in the early secretory pathway and to protein folding may contribute, in part, to improved laccase activity from these strains.

## Discussion

In this study, we performed a large-scale screen of laboratory, wild and industrial *S. cerevisiae* isolates by establishing a growth and screening pipeline for *ttLCC1* laccase expression using a library of ∼1,000 isolates. Following the identification of 47 candidate strains, we subsequently determined that 20 strains display a higher ability to produce active laccase with 4 strains producing higher ttLcc1 and mtLcc1 laccase activity. To uncover the biological factors that may contribute to improved activity by these strains, we characterized them using complementary genomic and proteomic approaches. Succeeding these analyses, we generated knockouts and confirmed improvement in laccase production in 10 of the deletions. Most of the knockouts that led to higher laccase activity were identified by determining genes significantly depleted among the isolated strains (8 out of 15 tested genes). In contrast, fewer validated targets were derived from the proteomic approach (2 out of 12 candidates). Intriguingly, we observed that many proteins are expressed at significantly different levels among the characterized strains, which may be exacerbated by the fact we grew the cell to saturation. One possibility is that the proteomic approach is currently limited by the lower number of strains characterized, which did not allow us to narrow down fewer candidate genes. Many genes and proteins affected in the 20 hit strains are involved in metabolism. It has been previously demonstrated that largescale metabolic re-wiring occurs in high α-amylase secreting mutant strains [30] and down regulation of genes involved in cellular metabolism can improve recombinant protein expression [22], several of our findings suggest a similar phenomenon may impact laccase production. A leading GO term among depleted genes is hexose catabolism driven by *HXT6-like*, *HXT11*, *IMA3* and *IMA5-like*. *HXT11* is a broad-spectrum hexose transporter [41] with roles in drug resistance [42] while *HXT6* is a glucose transporter [43] and the paralog of *HXT1*, the glucose transporter identified to be down regulated in mutant strains by Huang and colleagues [30]. In our assay, the *hxt11*Δ strain had an 85% increase in laccase activity. As Hxt11 is not required for glucose uptake, deletion of *HXT11* may lead to a change of allocated cellular resources and/or a reduced unnecessary glucose uptake during growth that could then favour heterologous protein production. Ima3 and Ima5 are both α-glucosidases with roles in isomaltose utilization. It has been demonstrated that there is high redundancy in glycolytic enzymes in *S. cerevisiae,* where deleting 13 of 27 minor paralogs did not result in any growth defects [44] even though glycolytic enzymes comprise a large portion of soluble proteins [45]. Neither *IMA3* nor *IMA5* are required for isomaltose utilization, so, similarly to *HXT11*, it is possible that downregulation of these genes silences the expression of these proteins which allows for allocation of resources towards recombinant protein production.

We provide further evidence that down regulation of thiamine biosynthesis pathways is beneficial to recombinant protein production, another point of agreement with Huang *et. al.* [30]. Both Sno2 and Snz3 are involved in biosynthesis of pyridoxine and subsequently thiamine, thiamine then feeds back and represses transcription of *SNZ3* [46, 47]. It is likely that *SNO2* is also repressed by thiamine as *SNZ* and *SNO* genes are co-regulated [48]. Once again, metabolic re-wiring may underlie the mechanism of how Snz3 and Sno2 depletion may contribute to improved laccase expression. Derivatives of both pyridoxine and thiamine are co-factors of numerous metabolic enzymes [49, 50]. Thus, down regulation of genes involved in their biosynthesis may lead to decreased levels of these co-factors in the cell, resulting in attenuation of the activity of metabolic enzymes.

An interesting group of depleted genes we identified among the 20 hit strains are the “compensated” genes, where homologous genes are also enriched. These included *GAL1, GAL7, GAL10, KAP104, PMU1 and PRM9* (Fig. 4B). We anticipated that the knockouts of these “compensated” genes in BY4741 would have no effect or be potentially detrimental to laccase production. Largely this was the case. *PMU1* deletion, which caused a greater decrease of laccase activity, is known to cause greater chromosome instability [51] and a defect in vacuole fragmentation [52], both likely contributing to decreased recombinant production. The *GAL* genes are all involved in galactose metabolism, a carbon source that is often omitted in laboratory settings, like in the media used in this study. Deletion of two of the three *GAL* genes tested (*gal1*Δ and *gal10*Δ) resulted in reduced laccase production (Fig. 4F). A *gal1*Δ strain has previously been noted to cause a decrease in endocytosis [53], a change in vacuolar morphology [52] and to increase the activity of an enzyme [54]. Similarly, *gal10*Δ has also been implicated in a decrease in endocytosis [55] and changes to vacuolar morphology [52]. Therefore, the negative impact on laccase production upon deletion of *GAL* genes could be due to a decrease in endocytosis, reducing their ability to acquire nutrients, and/or defects in vacuolar fragmentation reducing their ability to scavenge nutrients through microautophagy [56]. Contrary to the deletion of the *GAL* genes, *KAP104* and *PRM9* deletion strains both resulted in greater laccase production (Fig. 4F). Kap104 is a transportin involved in returning mRNA binding proteins to the nucleus [57] and deletion of *KAP104* causes mislocalization of Hrp1 and the mRNA binding protein Nab2 [58]. It is unknown what the mechanism behind increased laccase production in *kap104*Δ cells is, but possible explanations, among others, may be increased amino acids available for translation of new proteins by reducing levels of some mRNA. Prm9 is a member of the DUP240 family of membrane proteins with a COPII binding motif and is proposed to be involved in ER exit of proteins as Prm9 overexpression caused enlargement of the vacuole; however, Prm9 is not required for ER exit [38, 59]. Thus, Prm9 may play a role in targeting proteins for vacuolar degradation and deleting *PRM9* may redirect proteins to be secreted rather than be degraded in the vacuole. It is possible that *KAP104* and *PRM9* may not be faithfully compensated by their respective homologous genes in the identified strains (or simply have different functions), allowing their absence to contribute to the increased laccase production phenotype.

*PRM8* deletion, like its paralog *PRM9*, also resulted in significant increase to laccase production at a nearly identical level (Fig. S6 but I think should be Fig. 6G). Prm8 was detected to be significantly depleted in our proteomics studies (Additional file 2). Prm8 and 9 form a complex with each other at the ER membrane [38] and it is likely that the effect of deleting one of the paralogs impairs the function of the complex, thus, accounting for their similar effects on laccase production. Like *prm9*Δ*, prm8*Δ may impair vacuolar targeting of laccase, redirecting it to be secreted from the cell instead.

We expected chaperones and other protein quality control proteins to play a larger role in impacting recombinant protein production. However, we failed to identify many changes to the levels of chaperones in the protein mass spectrometry experiment, apart from a notable reduction of *SSE2* and its paralog *SSE1* in many of the characterized strains. While deletion of *SSE2* did not result in significant improvement in laccase activity, deletion of *SSE1* did (Fig. 6F). Whereas both *SSE1* and *SSE2* are upregulated under heat stress conditions, Sse1 is present at higher levels than Sse2 in normal growth temperatures [60, 61], potentially explaining the different impact of the deletions of these two genes on laccase activity. More work would be required to determine how deletion of Sse1, a nucleotide exchange factor for cytosolic Hsp70s, may impact laccase activity.

In contrast, we found that many vacuolar proteins were expressed at lower levels in the proteomic experiment. We previously identified S288C gene deletions causing defects in vacuolar protein sorting, (including numerous *VPS* gene deletions), that improved heterologous laccase expression, indicating that the vacuole is a key player in recombinant protein secretion [21]. Notably, one of the common targets for strain engineering to increase secreted recombinant protein yields is the deletion of *PEP4*, also a vacuolar protease. We detected lower levels of three vacuolar enzymes (the phospholipase Atg15, the peptidase Cps1 and the putative protease Ysp3), known or suspected to be reliant on Pep4 to mature and targeted to the vacuole *via* the multivesicular body pathway [62–64]. Notably, Atg15 is essential for breakdown of autophagic membranes in the vacuole to release the contents for degradation [34, 65] that is supported by enzymes like Cps1 and Ysp3 [66, 67]. While our work suggests that modulating vacuolar degradation could positively impact laccase production, none of the assessed single deletion genes associated with vacuolar degradation increased laccase activity (Fig. 6E). One possibility is that multiple vacuolar genes need to be targeted, or the levels of these proteins need to be carefully tuned in to have an impact on recombinant protein production.

## Methods

### Plasmid and Yeast Strain Construction

All plasmids, oligos and yeast strains utilized in this study are listed in Additional file 1. To construct the *ttLCC1* laccase expression plasmid (BPM1747), the *URA3* cassette from pRS416-GPD1p-*ttLCC1*-CYC1t, a gift from Dr. Sychrová [23], was swapped with the *kanMX6* cassette from pFA6a-*kanMX6* [68] using Gibson assembly. The same approach was used to generate the control empty vector (BPM1743) using pRS416 [69]. The *mtLCC1* gene was subcloned with the *GPD1* promoter and *CYC1* terminator from pVT100U-MtL, also a gift from Dr. Sychrová, into the same *kanMX6* containing vector (to generate BPM1752) using Gibson assembly. This *mtLCC1* is the T2 variant generated by Butler *et. al.* [14].

The yeast strain library was de-condensed from its original 384-format using an S&P Robotics BM3-BC into a 96-format for strain recovery and long-term storage. Subsequent handling of the de-condensed library was done with a manual pinning tool. The library was transformed with BPM1747 in 96-well format *via* the high-throughput LiAc transformation protocol [70], allowing development of drug resistance with 3 hours of recovery in YPD before selection. Successful transformants were selected for by plating on YPD + G418 (200 µg/mL).

Single gene deletion strains were generated by homologous recombination integrating a *HIS3* cassette from pFA6a-*His3MX6* at the target locus. Knockout cassettes containing 60bp of homology to the target locus on either side were generated by PCR amplification and transformed into BY4741 *via* the high-efficiency LiAc/ssDNA method [71]. Gene deletions were verified by colony PCR confirming integration of the cassette at the target locus and absence of the target gene.

### ABTS-based Laccase Activity Assay and Quantification

10 colonies were pooled together per replicate of each transformed strain (individual colonies were tested for the *HIS3* integration controls) and precultured in 150 µL of YPD + G418 (200 µg/mL) overnight at 30°C in a 96-well round bottom plate with shaking at 900 rpm. The precultures were used to inoculate 1 mL of laccase expression medium (YPD, 20 μg/mL adenine, 50 mM potassium phosphate (dibasic, pH 6), 0.5 mM copper (II) sulfate) with 1/2× G418 (100 µg/mL) to a starting OD_600_ of 0.2 in a 2 mL 96-deep-well round bottom plate. The laccase expression cultures were grown for 4 days (96 h) at 30°C with shaking at 900 rpm. At the end of the 4 days, the final OD_600_ of each culture was measured before the cells were pelleted by centrifugation at 3,200 rcf for 5 min. 100 µL of the cleared media containing the secreted laccase was transferred to a clear-96-well flat bottom plate to be used for laccase activity assays. Immediately before quantitation, 100 µL of 2 mM ABTS in 100 mM Britton and Robinson buffer (100 mM each of boric, phosphoric and acetic acid, brought to pH 4 by addition of NaOH) was added to the cleared media to begin the reaction [72]. Laccase activity was monitored by UV-visual spectrometry (absorbance) at 420 nm over the course of 1 hour with readings every minute starting from minute 0, in the Clariostar + plate reader (BMG). We used double orbital shaking at 300 RPM for 30s between reads, and centre point reading with 20 flashes. Ab_420nm_ was plotted against time and linear regression curves were fitted to the data (data points on both ends of the curve were removed until an R^2^ of at least 0.999 was achieved to only take in account the linear range of the reaction). The Beer-Lambert law was used to convert absorbance into concentration of oxidized ABTS in µmols using a molar extinction coefficient of 36,000 M^−1^ cm^−1^ [73]. This was then used to calculate laccase activity (1 unit of laccase activity = 1 µmol oxidized ABTS / min).

### Phylogenetic Tree

A neighbour-joining phylogenetic tree was generated and exported in Newick format using the “ape” package in R and the “1011distanceMatrixBasedOnSNPs.tab” dataset from Peter, J. *et. al.* [25]. The clades were annotated, and strains highlighted on the phylogenetic tree using the Interactive Tree of Life (iTOL) webtool [74]. The tree was rooted to the strain HN6 of the CHN I clade, due to the suspected origin of *S. cerevisiae* from China [25, 75] and high genetic diversity of strains from this origin [76].

### RT-qPCR

For RT-qPCR experiments, total RNA was extracted using the RiboPure Yeast RNA Prep Kit (Thermo Fisher Scientific AM1926) from an equivalent of 25 ODs of cells from 4 biological replicates. A Nanodrop One (Thermo Fisher Scientific) was used to determine the crude RNA concentration for dilution of the RNA preps into a suitable range for precise RNA concentration and RNA integrity number (RIN) determination using a Bioanalyzer 2100 (Agilent G2939BA) and RNA 6000 Nano chip (Agilent 5067−1511). All samples displayed a RIN of 6.8 or more (Additional file 3), above the recommended RIN for RT-qPCR [77, 78]. *ttLCC1* mRNA levels were determined using the Power SYBR® Green RNA-to-CT™ 1-Step Kit (Thermo Fisher Scientific 4389986) using 20 ng of RNA and 10 μM primers (Additional file 1) per replicate. *UBC6* was used as the internal reference gene. Three 20 μL replicates were pipetted into a 384-well PCR plate for each strain and target, the plate was sealed with an optically clear seal and the RT-qPCR was run in a ViiA 7 Real-Time PCR System (Thermo Fisher Scientific 4453545) with cycle settings following manufacturer’s protocols for the RT-qPCR kit. Data was visualized and exported to Excel using QuantStudio Real-Time PCR Software (Thermo Fisher Scientific v1.6.1). The comparative *C_T_* (ΔΔ*C_T_*) method was used to determine the relative *ttLCC1* mRNA levels [79].

### GWAS

Mixed-model association analysis was performed using the FaST-LMM python library version 0.2.32 (https://fastlmm.github.io/ and https://pypi.org/project/fastlmm/0.2.32/) [80]. We used the normalized phenotypes by replacing the observed value by the corresponding quantile from a standard normal distribution, as FaST-LMM expects normally distributed phenotypes. The command used for association testing was the following: single_snp(bedFiles, pheno_fn, count_A1 = True), where bedFiles is the path to the PLINK formatted SNP data and pheno_fn is the PLINK formatted phenotype file.

### Statistical Calculations and Gene Ontology Enrichment Analysis

Statical calculations were done in R (Rstudio 2023.06.1+524, R version 4.3.3) using custom scripts and the “stats” package of R (https://github.com/RyanWKW/Mining-Yeast-Diversity-Unveils-Novel-Targets-for-Improved-Heterologous-Laccase-Production-in-Sacchar) and all p-values are listed in Additional file 4. Statistical enrichment of ORFs was performed using Fisher’s exact test to compare the proportional presence of ORFs in the hits to the remainder of the library. Only genes that were either absent or present in the 20 hits strains in comparison to the reference strain were first retained. We then compared their proportional presence/absence in the 20 strains to their proportional presence/absence in the remaining 1000+ strains on the library using the Fisher’s exact. Fisher’s exact tests were done using the “fisher.test” function. T-tests for the mass spectrometry analysis and the laccase activity assays were conducted using the “t.test” function and corrected, if necessary, using the Benjamini-Hochberg method with the “p.adjust” function. Pearson correlation was calculated using the “cor.test” function. Median absolute deviations (MAD) were calculated using the formula MAD = 1.4826* Median (|X_i_ - median(X)|). The proteome correlation heatmap was generated by comparing the data using a linear model with the “lm” function. GO term enrichment analysis was performed using the online tool, ShinyGO 0.80 [81], against the *Saccharomyces cerevisiae* background and using an FDR cut-off of 0.05. The split axis in Figure 3B was created with the “ggbreak” add-in for ggplot2 [82].

### Proteomics

Pelleted cells from 1 mL expression cultures were washed 3 times with 500 µL of 50 mM TRIS-HCl (pH6.8) before being lysed in 50 mM TRIS-HCl (pH 6.8) with 2% SDS using a Precellys 24 (Bertin) and 400 µm acid washed silica beads (OPS Diagnostics BAWG 400-200-04). 5 µg of proteins from the lysates were digested following the SP3 protocol [83] with Sera-Mag™ carboxylate-modified SpeedBeads [E7] and [E3] (Cytiva 45152105050250 and 65152105050250) and sequencing grade modified trypsin (Promega V5113). The digested peptides were acidified with trifluoroacetic acid (TFA) before desalting. Peptides were desalted using an AssayMAP Bravo (Agilent) liquid handler and AssayMAP 5 µL C18 cartridges (Agilent 5190-6532). Cartridges were primed with 100 µL of priming buffer (0.1% TFA, 80% acetonitrile (AcN)) then washed with 50 µL of buffer A (0.1% TFA, 5% AcN) before loading of the peptides. A “cup wash” step was performed with 25 µL of buffer A, then a sample wash was done with 50 µL of buffer A. The peptides were eluted from the C18 with buffer B (0.1% TFA, 40% AcN) before drying with a Vacufuge plus (Eppendorf 022820109). Dried peptides were reconstituted in 0.1% TFA & 0.5% AcN before loading on a timsTOF Pro 2 operated in DIA-PASEF mode coupled to a NanoElute UHPLC system (Bruker Daltonics). For each sample, 50 ng was loaded on an Aurora Series Gen2 analytical column heated to 50 °C. The analytical column was equilibrated with buffer A (0.1% FA and 0.5% AcN in water) then subjected to a 30-min gradient with a 0.3 µL/min flow. Buffer B (0.1% FA, 0.5% water in AcN) was increased from 2 to 12% over the first 15 min then to 33% from 15 to 30 min, followed by 95% over 30 s and held at 95% for 7.72 min. The DIA acquisition scanning ranged from 100 to 1,700 m/z. Following the MS1 scan, 17 PASEF scans of 22, 35 m/z windows ranging from 319.5 to 1,089.5 m/z were performed. Ion mobilities ranged from 0.7 to 1.35 V s/cm2 with a 100-ms ramp and accumulation time and a 9.42-Hz ramp rate resulting in a 1.91-s cycle time. Collision energy was increased linearly as a function of ion mobility from 27eV at 1/k0 = 0.7 V s/cm2 to 55 eV at 1/k0 = 1.35 V s/cm2.

Data files were searched using the directDIA pipeline of Spectronaut (Biognosys ver. 18.7.240506.55695), quantifying proteins using the top 3 peptides at the MS2 level. Quality control on the data was done by calculating the CVs and determining correlation of the data sets using a custom script fitting to a linear model using the “lm” function of the “stats” package in R. Principal component analysis was conducted using the “PCA” function of the “FactoMineR” package in R. Strain comparison plots were created with “ggplot2”, using averaged intensities of proteins quantified in both strains (https://github.com/RyanWKW/Mining-Yeast-Diversity-Unveils-Novel-Targets-for-Improved-Heterologous-Laccase-Production-in-Sacchar). Before running t-tests, the protein quantities were normalized across all datasets by first dividing each quantity by the median within the dataset, then multiplying by the mean of median quantities across datasets, and missing values were imputed by random assignment of the bottom 5% of intensities using the “impute.MinProb” function of the “imputeLCMD” package of R, tuning the sigma to ensure no negative values are imputed.

## Supporting information

Additonal file 1

## Acknowledgements

The pRS416 plasmid was gifted by Dr. Philip Hieter, and the *ttLCC1* and *mtLCC1* with promoter and terminator constructs were gifted by Dr. Hana Sychrová. We thank Jason Rogalski, Renata Moravcová, Jeanne Yuan and Lok Tin Hui for mass spectrometry instrument maintenance and sample loading, Michelle Moksa and Qi Cao from the Hirst Lab for their assistance with RT-qPCR experiments, Uche Joseph Ogbede from the Nislow lab for technical help, Renaissance Biosciences for providing some yeast strains and all members of the Mayor lab for their input and discussion. Figure 1A was created with BioRender.com. This work is supported by a NSERC Discovery Grant (RGPIN-2022-03787) and a CFI Innovation Fund (39914).

## Author Contributions

R.W. designed, executed, and analyzed the data of most experiments. M.F. generated some plasmids and assisted in establishment of the screening pipeline. J.L. generated and screened some single gene deletions. T.W. assisted in secondary laccase screens. J.M. generated and annotated the phenotypic tree. F.D. performed the GWAS analysis. C.M. created the script for correlation heatmapping. V.M. and J.S. provided yeast strains critical to the work. C.N. provided access to critical infrastructure. T.M. helped design the experiments and analyze the data. The manuscript was written by R.W. and T.M. and edited by all the authors.

## Data availability

All data generated or analysed during this study are included in this published article and its supplementary information files. Mass Spectrometry files are available on MassIVE (XXX). Additional datasets generated during the current study are available from the corresponding author upon reasonable request. Plasmids and yeast strains listed in the supplemental material with a BPM or YTM denomination are available upon request.

---

**Fig. S1.**
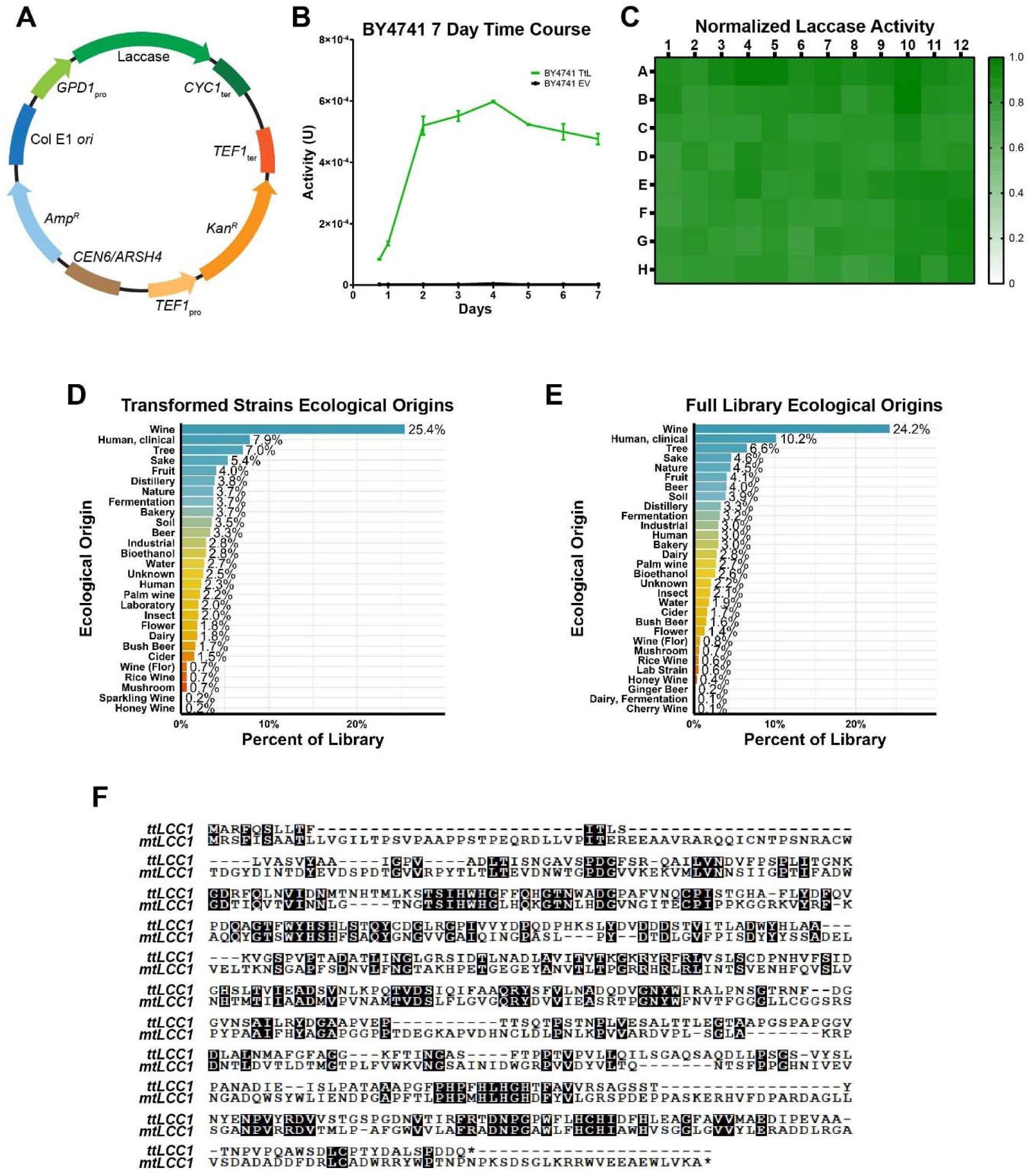
**A)** Plasmid map of BPM1747 with laccase. **B)** Laccase activity from BY4741 cells expressing and secreting *ttLCC1* was measured over a 7-day period. **C)** Heatmap plotting relative laccase activity from BY4741 cells expressing *ttLCC1* for 4-days in a 96-deep-well plate. **D and E)** Ecological origin distributions of the full library (D) and of the 597 transformed strains from which the laccase activity was assessed (E). **F)** Protein sequence alignment of *ttLCC1* and *mtLCC1* highlighting sequence identity.

**Fig. S2.**
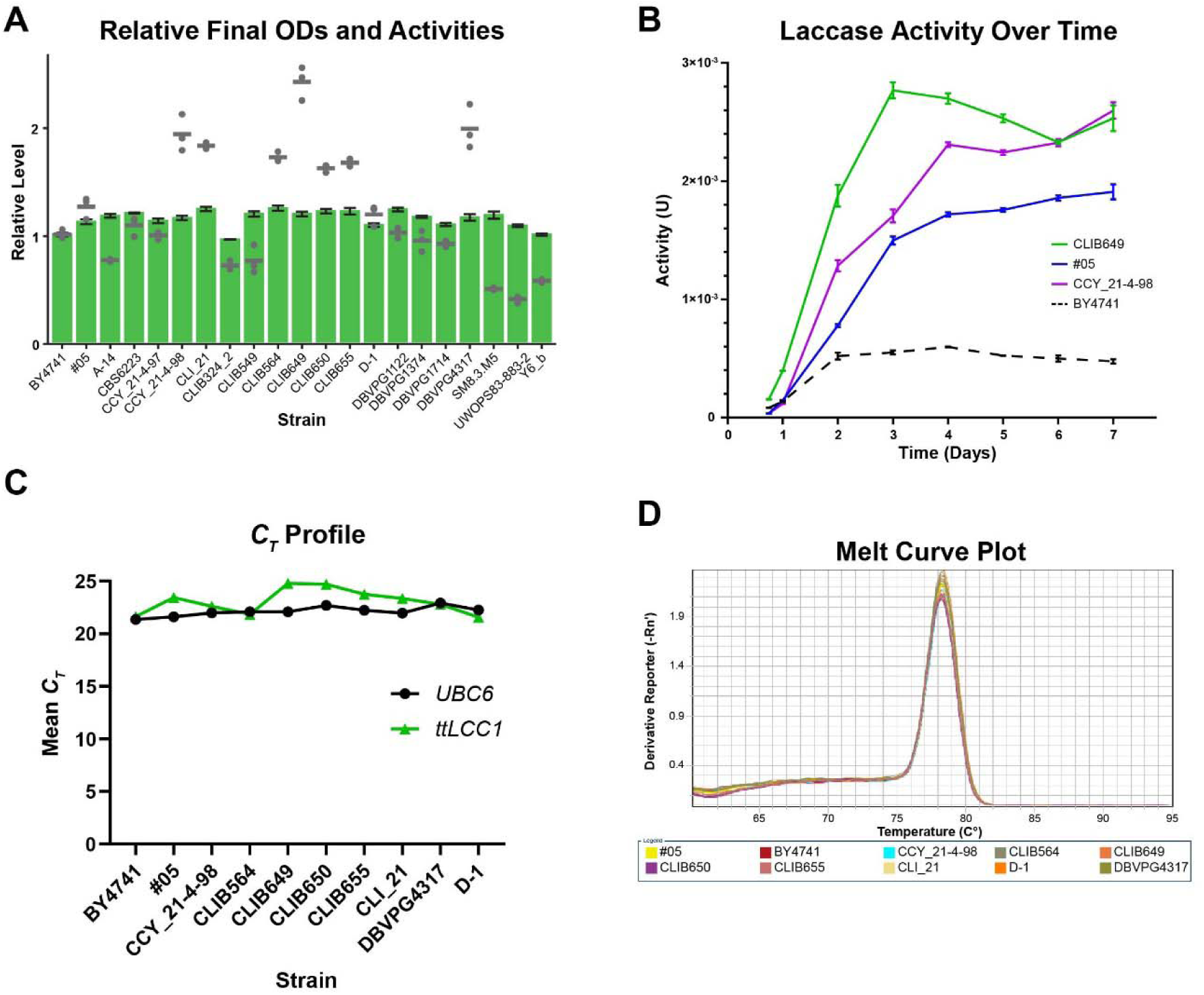
**A)** Relative cell density (represented by bars) and laccase activity (points) in the indicated strains expressing *ttLCC1* after 4-days growth in comparison to BY4741 for the experiment shown in Fig. 1C. **B)** Laccase activity for 3 of the hit strains grown in a 25mL culture was assessed over 7-days. Activity from BY4741 cells shown in Fig. S1B was replotted for comparison. **C)** *C_T_* profile of the internal reference *UBC6* and target *ttLCC1* mRNA in the indicated strains. **D)** Melting curve of *ttLCC1* target demonstrates a single target was amplified.

**Fig. S3.**
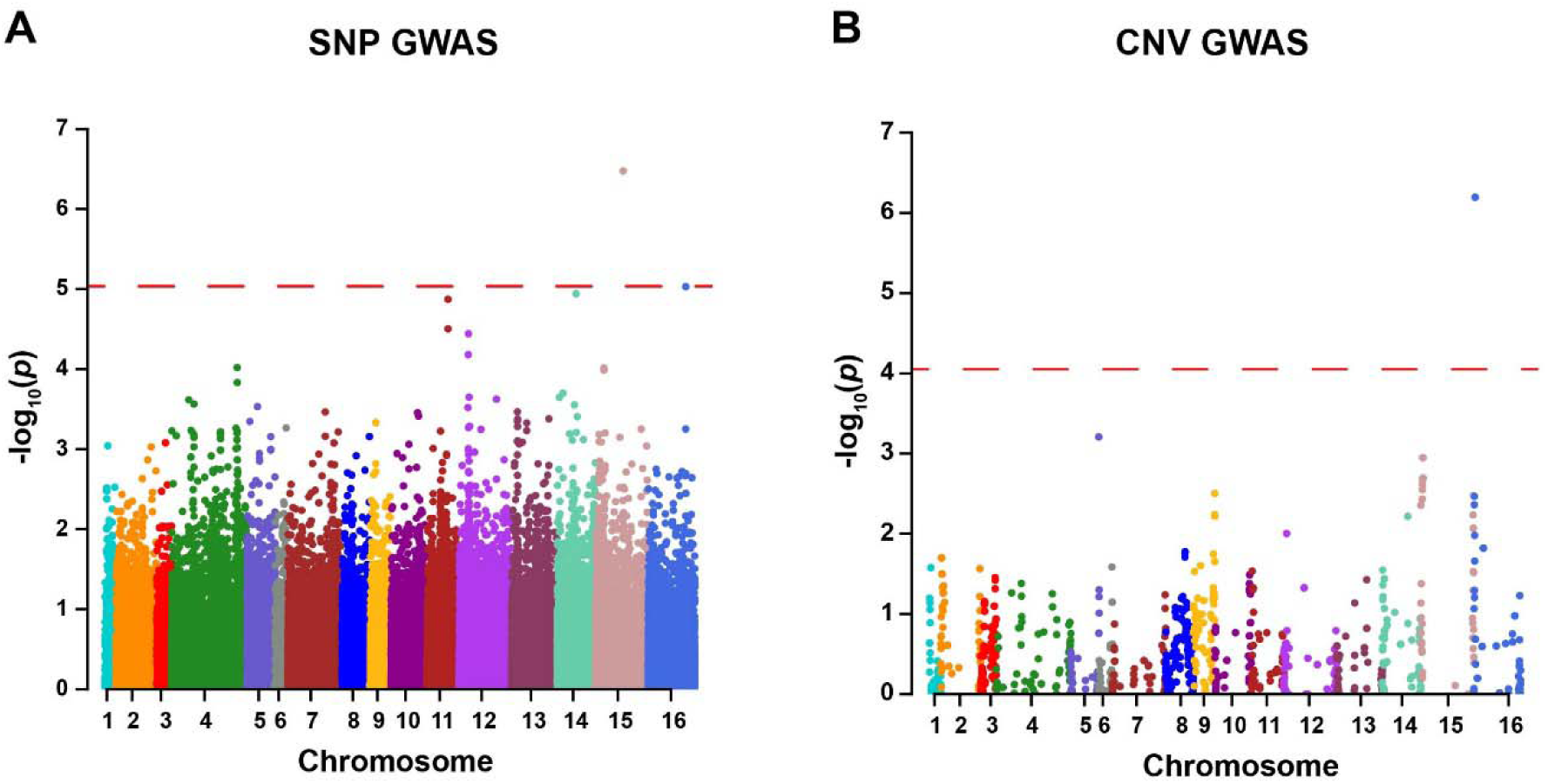
**A)** SNP GWAS identified a single significant SNP in *ATG27/ETF1* of chromosome 15. **B)** CNV GWAS identified a single significant CNV in the locus encompassing YPL273W and YPL274W on chromosome 16.

**Fig S4.**
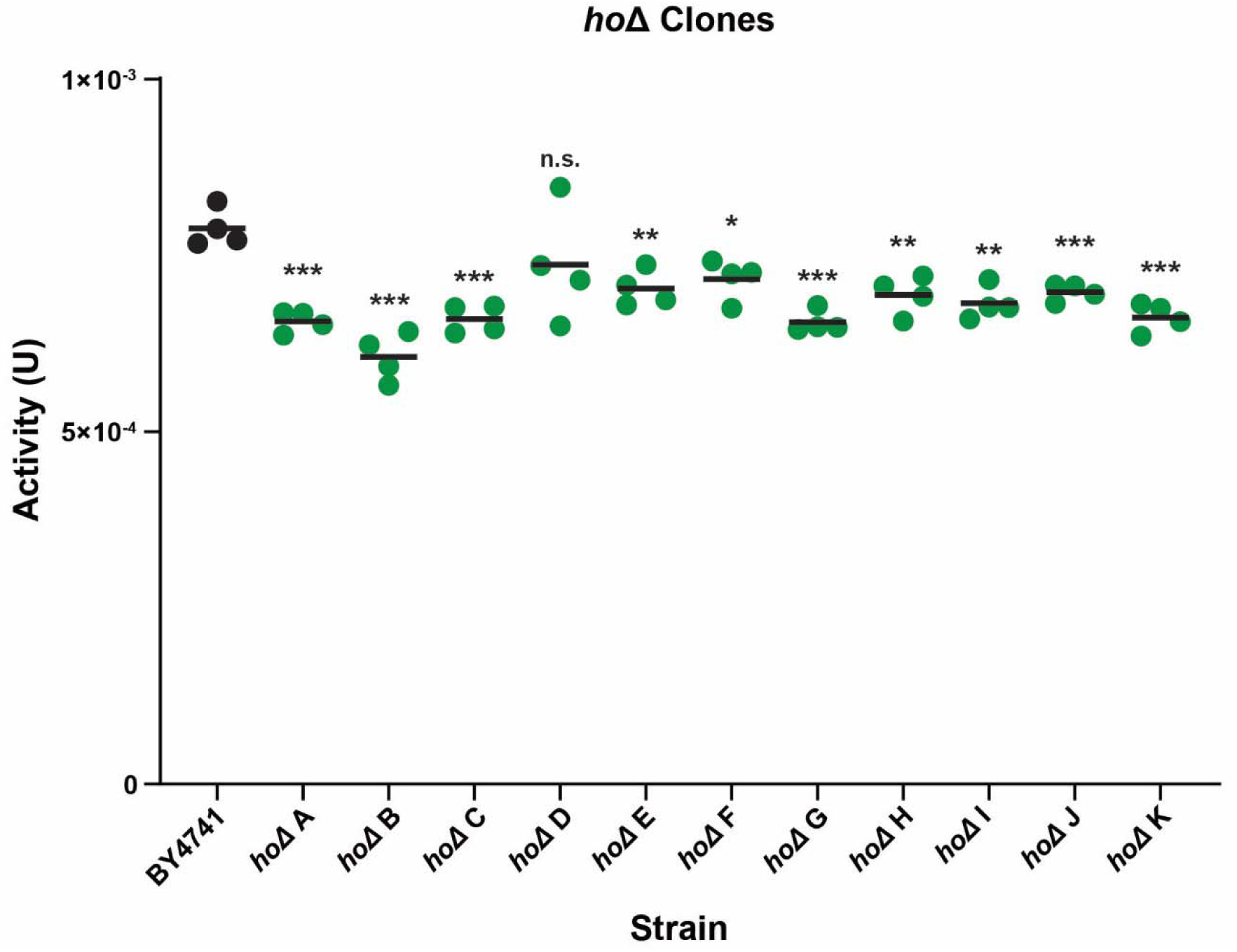
Laccase activity of individual clones after 4 days growth (n=4; p-values: * < 0.05, ** < 0.01 *** < 0.001, **** < 0.0001).

**Fig. S5.**
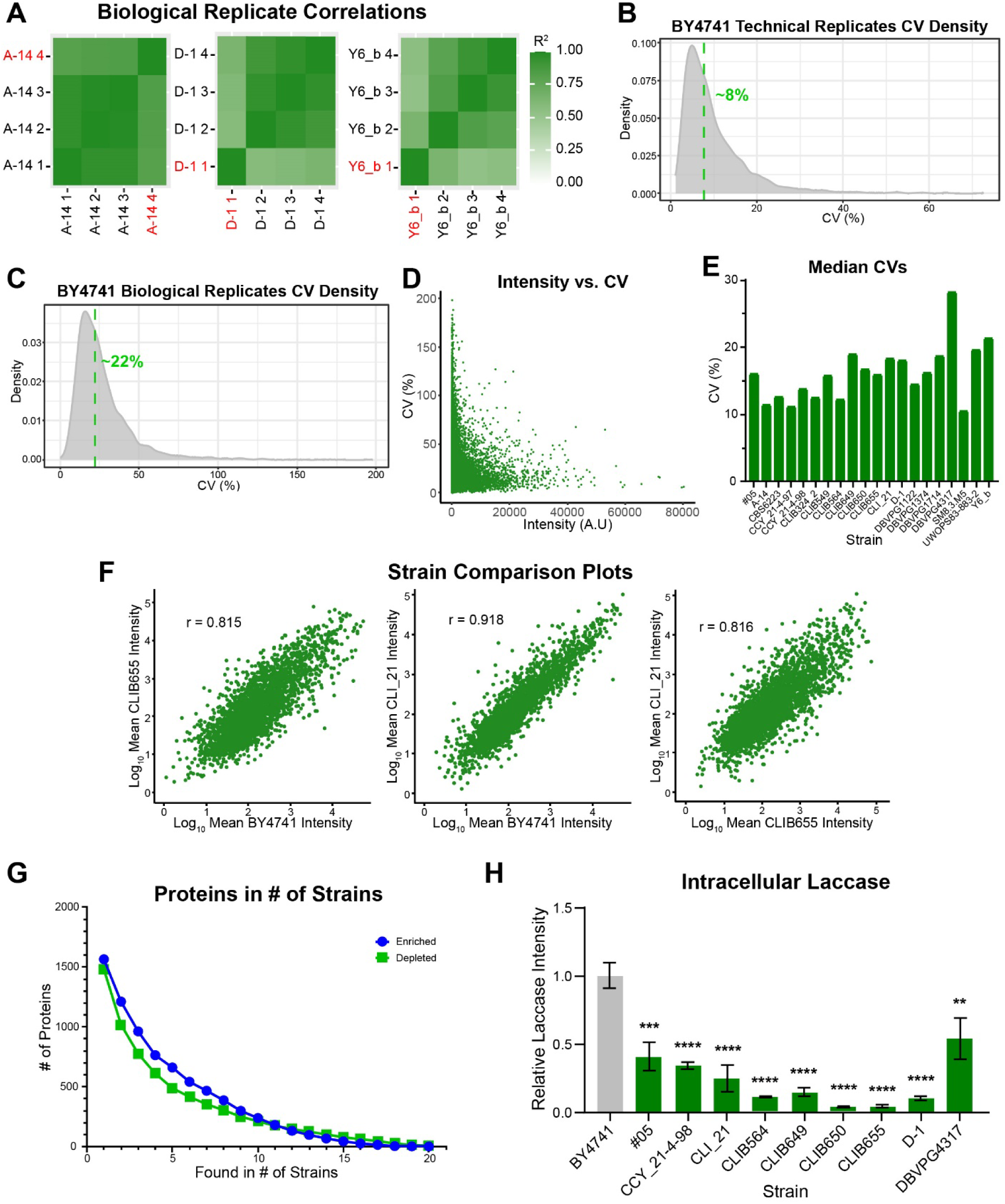
**A)** Correlation heatmaps of strains A-14, D-1 and Y6_b. Replicates indicated in red were removed from further analysis due to low correlations indicative of potential errors during sample preparation. **B and C)** CV density plots of the technical replicates of one BY4741 sample (n=14) and biological replicates (n=7) of BY4741 cells grown in different wells. Median CVs shown (green dotted lines). **D)** Scatter plot of CVs of each protein identified in each mass spectrometry run of the 20 hit strains and BY4741 against its averaged intensity. CVs steeply increase for some proteins with low intensities. **E)** Plot of the median CVs of the 20 hit strains. **F)** Correlation plots of selected strains. Only proteins quantified in both strains are shown after averaging their intensities from the biological replicates. Pearson correlation indicated. **G)** The number of significantly enriched and depleted proteins in the indicated number of strains in comparison to BY4741. 1565 proteins are found to be enriched in at least 1 of the strains, down to a single protein enriched in all 20 strains. Similarly, 1481 proteins are depleted in at least 1 of the strains and 8 proteins are depleted in all 20. **H)** Relative intracellular ttLcc1 intensities of the indicated strains after 4-days growth (BY4741: n=7; D-1: n = 3; remaining: n=4; p-values: * < 0.05, ** < 0.01 *** < 0.001, **** < 0.0001).

**Fig. S6.**
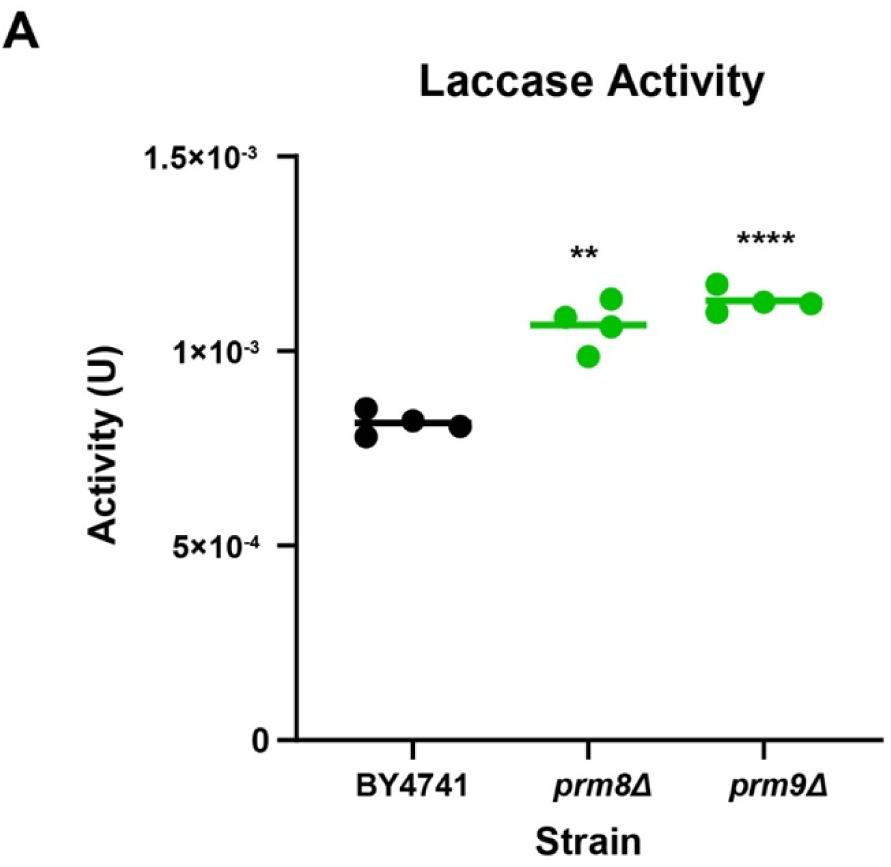
Laccase activity in the indicated strains expressing *ttLCC1* after 4-days growth (n=4; p-values: * < 0.05, ** < 0.01 *** < 0.001, **** < 0.0001).

## Additional Files

Filename: Additional file 1

Format: Excel worksheet (.xlsx)

Title: Plasmids, Oligos and Strains

Description: Tables of all plasmids, oligos and strains used or generated in this work.

Filename: Additional file 2

Format: Excel worksheet (.xlsx)

Title: Proteins Quantified

Description: Tables of all proteins quantified in this work. Includes the normalized and outlier removed dataset.

Filename: Additional file 3

Format: Excel worksheet (.xlsx)

Title: RIN

Description: RNA integrity numbers of each RNA prep used for RT-qPCR.

Filename: Additional file 4

Format: Excel worksheet (.xlsx)

Title: P-values

Description: Tables of all p-values from statistical calculations in this work.

## References

1. Wang G, Huang M, Nielsen J. Exploring the potential of Saccharomyces cerevisiae for biopharmaceutical protein production. Curr Opin Biotechnol. 2017;48:77–84.

2. Arregui L, Ayala M, Gómez-Gil X, Gutiérrez-Soto G, Hernández-Luna CE, Herrera de Los Santos M, et al. Laccases: structure, function, and potential application in water bioremediation. Microb Cell Fact. 2019;18:200.

3. Messerschmidt A. Blue Copper Oxidases. 1993. p. 121–85.

4. Sugumaran M, Giglio L, Kundzicz H, Saul S, Semensi V. Studies on the enzymes involved in puparial cuticle sclerotization in Drosophila melanogaster. Arch Insect Biochem Physiol. 1992;19:271–83.

5. Andersen SO. Insect cuticular sclerotization: a review. Insect Biochem Mol Biol. 2010;40:166–78.

6. Givaudan A, Effosse A, Faure D, Potier P, Bouillant M-L, Bally R. Polyphenol oxidase in Azospirillum lipoferum isolated from rice rhizosphere: Evidence for laccase activity in non-motile strains of Azospirillum lipoferum. FEMS Microbiol Lett. 1993;108:205–10.

7. Faure D, Bouillant ML, Bally R. Isolation of Azospirillum lipoferum 4T Tn5 Mutants Affected in Melanization and Laccase Activity. Appl Environ Microbiol. 1994;60:3413–5.

8. Berrocal MM, Rodríguez J, Ball AS, Pérez-Leblic MI, Arias ME. Solubilisation and mineralisation of [14C]lignocellulose from wheat straw by Streptomyces cyaneus CECT 3335 during growth in solid-state fermentation. Appl Microbiol Biotechnol. 1997;48:379–84.

9. Bourbonnais R, Paice MG, Reid ID, Lanthier P, Yaguchi M. Lignin oxidation by laccase isozymes from Trametes versicolor and role of the mediator 2,2’-azinobis(3-ethylbenzthiazoline-6-sulfonate) in kraft lignin depolymerization. Appl Environ Microbiol. 1995;61:1876–80.

10. Eggert C, Temp U, Dean JFD, Eriksson K-EE. A fungal metabolite mediates degradation of non-phenolic lignin structures and synthetic lignin by laccase. FEBS Lett. 1996;391:144–8.

11. Garzillo AM, Colao MC, Caruso C, Caporale C, Celletti D, Buonocore V. Laccase from the white-rot fungus Trametes trogii. Appl Microbiol Biotechnol. 1998;49:545–51.

12. Youn H-D, Hah YC, Kang S-O. Role of laccase in lignin degradation by white-rot fungi. FEMS Microbiol Lett. 1995;132:183–8.

13. Mate DM, Alcalde M. Laccase: a multi-purpose biocatalyst at the forefront of biotechnology. Microb Biotechnol. 2017;10:1457–67.

14. Bulter T, Alcalde M, Sieber V, Meinhold P, Schlachtbauer C, Arnold FH. Functional expression of a fungal laccase in Saccharomyces cerevisiae by directed evolution. Appl Environ Microbiol. 2003;69:987–95.

15. Shusta E V., Raines RT, Plückthun A, Wittrup KD. Increasing the secretory capacity of Saccharomyces cerevisiae for production of single-chain antibody fragments. Nat Biotechnol. 1998;16:773–7.

16. Hackel BJ, Huang D, Bubolz JC, Wang XX, Shusta E V. Production of soluble and active transferrin receptor-targeting single-chain antibody using Saccharomyces cerevisiae. Pharm Res. 2006;23:790–7.

17. Tang H, Bao X, Shen Y, Song M, Wang S, Wang C, et al. Engineering protein folding and translocation improves heterologous protein secretion in Saccharomyces cerevisiae. Biotechnol Bioeng. 2015;112:1872–82.

18. Jones EW. Tackling the protease problem in Saccharomyces cerevisiae. Methods Enzymol. 1991;194 C:428–53.

19. Tomimoto K, Fujita Y, Iwaki T, Chiba Y, Jigami Y, Nakayama K, et al. Protease-deficient Saccharomyces cerevisiae strains for the synthesis of human-compatible glycoproteins. Biosci Biotechnol Biochem. 2013;77:2461–6.

20. Huang M, Bai Y, Sjostrom SL, Hallström BM, Liu Z, Petranovic D, et al. Microfluidic screening and whole-genome sequencing identifies mutations associated with improved protein secretion by yeast. Proc Natl Acad Sci U S A. 2015;112:E4689–96.

21. Strawn G, Wong RWK, Young BP, Davey M, Nislow C, Conibear E, et al. Genome-wide screen identifies new set of genes for improved heterologous laccase expression in Saccharomyces cerevisiae. Microb Cell Fact. 2024;23:36.

22. Wang G, Björk SM, Huang M, Liu Q, Campbell K, Nielsen J, et al. RNAi expression tuning, microfluidic screening, and genome recombineering for improved protein production in Saccharomyces cerevisiae. Proc Natl Acad Sci U S A. 2019;116:9324–32.

23. Antošová Z, Herkommerová K, Pichová I, Sychrová H. Efficient secretion of three fungal laccases from Saccharomyces cerevisiae and their potential for decolorization of textile industry effluent-A comparative study. Biotechnol Prog. 2018;34:69–80.

24. Niku-Paavola ML, Raaska L, Itävaara M. Detection of white-rot fungi by a non-toxic stain. Mycol Res. 1990;94:27–31.

25. Peter J, De Chiara M, Friedrich A, Yue J-X, Pflieger D, Bergström A, et al. Genome evolution across 1,011 Saccharomyces cerevisiae isolates. Nature. 2018;556:339–44.

26. Kawai S, Hashimoto W, Murata K. Transformation of Saccharomyces cerevisiae and other fungi: methods and possible underlying mechanism. Bioeng Bugs. 2010;1:395–403.

27. Brachmann CB, Davies A, Cost GJ, Caputo E, Li J, Hieter P, et al. Designer deletion strains derived from Saccharomyces cerevisiae S288C: a useful set of strains and plasmids for PCR-mediated gene disruption and other applications. Yeast. 1998;14:115–32.

28. Ralser M, Kuhl H, Ralser M, Werber M, Lehrach H, Breitenbach M, et al. The Saccharomyces cerevisiae W303-K6001 cross-platform genome sequence: insights into ancestry and physiology of a laboratory mutt. Open Biol. 2012;2:120093.

29. Teste M-A, Duquenne M, François JM, Parrou J-L. Validation of reference genes for quantitative expression analysis by real-time RT-PCR in Saccharomyces cerevisiae. BMC Mol Biol. 2009;10:99.

30. Huang M, Bao J, Hallström BM, Petranovic D, Nielsen J. Efficient protein production by yeast requires global tuning of metabolism. Nat Commun. 2017;8:1131.

31. Muenzner J, Trébulle P, Agostini F, Zauber H, Messner CB, Steger M, et al. Natural proteome diversity links aneuploidy tolerance to protein turnover. Nature. 2024;630:149–57.

32. Teyssonnière EM, Trébulle P, Muenzner J, Loegler V, Ludwig D, Amari F, et al. Species-wide quantitative transcriptomes and proteomes reveal distinct genetic control of gene expression variation in yeast. Proc Natl Acad Sci U S A. 2024;121:e2319211121.

33. Warringer J, Zörgö E, Cubillos FA, Zia A, Gjuvsland A, Simpson JT, et al. Trait variation in yeast is defined by population history. PLoS Genet. 2011;7:e1002111.

34. Epple UD, Suriapranata I, Eskelinen EL, Thumm M. Aut5/Cvt17p, a putative lipase essential for disintegration of autophagic bodies inside the vacuole. J Bacteriol. 2001;183:5942–55.

35. Lang T, Reiche S, Straub M, Bredschneider M, Thumm M. Autophagy and the cvt pathway both depend on AUT9. J Bacteriol. 2000;182:2125–33.

36. Davey HM, Cross EJM, Davey CL, Gkargkas K, Delneri D, Hoyle DC, et al. Genome-wide analysis of longevity in nutrient-deprived Saccharomyces cerevisiae reveals importance of recycling in maintaining cell viability. Environ Microbiol. 2012;14:1249–60.

37. VanderSluis B, Hess DC, Pesyna C, Krumholz EW, Syed T, Szappanos B, et al. Broad metabolic sensitivity profiling of a prototrophic yeast deletion collection. Genome Biol. 2014;15:R64.

38. Sandmann T, Herrmann JM, Dengjel J, Schwarz H, Spang A. Suppression of coatomer mutants by a new protein family with COPI and COPII binding motifs in Saccharomyces cerevisiae. Mol Biol Cell. 2003;14:3097–113.

39. Dragovic Z, Broadley SA, Shomura Y, Bracher A, Hartl FU. Molecular chaperones of the Hsp110 family act as nucleotide exchange factors of Hsp70s. EMBO J. 2006;25:2519–28.

40. Liu XD, Morano KA, Thiele DJ. The yeast Hsp110 family member, Sse1, is an Hsp90 cochaperone. J Biol Chem. 1999;274:26654–60.

41. Jordan P, Choe J-Y, Boles E, Oreb M. Hxt13, Hxt15, Hxt16 and Hxt17 from Saccharomyces cerevisiae represent a novel type of polyol transporters. Sci Rep. 2016;6:23502.

42. Nourani A, Wesolowski-Louvel M, Delaveau T, Jacq C, Delahodde A. Multiple-drug-resistance phenomenon in the yeast Saccharomyces cerevisiae: involvement of two hexose transporters. Mol Cell Biol. 1997;17:5453–60.

43. Reifenberger E, Freidel K, Ciriacy M. Identification of novel HXT genes in Saccharomyces cerevisiae reveals the impact of individual hexose transporters on glycolytic flux. Mol Microbiol. 1995;16:157–67.

44. Solis-Escalante D, Kuijpers NGA, Barrajon-Simancas N, van den Broek M, Pronk JT, Daran J-M, et al. A Minimal Set of Glycolytic Genes Reveals Strong Redundancies in Saccharomyces cerevisiae Central Metabolism. Eukaryot Cell. 2015;14:804–16.

45. Herrero P, Galíndez J, Ruiz N, Martínez-Campa C, Moreno F. Transcriptional regulation of the Saccharomyces cerevisiae HXK1, HXK2 and GLK1 genes. Yeast. 1995;11:137–44.

46. Nosaka K. Recent progress in understanding thiamin biosynthesis and its genetic regulation in Saccharomyces cerevisiae. Appl Microbiol Biotechnol. 2006;72:30–40.

47. Paxhia MD, Downs DM. SNZ3 Encodes a PLP Synthase Involved in Thiamine Synthesis in Saccharomyces cerevisiae. G3 (Bethesda). 2019;9:335–44.

48. Padilla PA, Fuge EK, Crawford ME, Errett A, Werner-Washburne M. The highly conserved, coregulated SNO and SNZ gene families in Saccharomyces cerevisiae respond to nutrient limitation. J Bacteriol. 1998;180:5718–26.

49. Parra M, Stahl S, Hellmann H. Vitamin B and Its Role in Cell Metabolism and Physiology. Cells. 2018;7:84.

50. Hohmann S, Meacock PA. Thiamin metabolism and thiamin diphosphate-dependent enzymes in the yeast Saccharomyces cerevisiae: genetic regulation. Biochim Biophys Acta. 1998;1385:201–19.

51. Choy JS, O’Toole E, Schuster BM, Crisp MJ, Karpova TS, McNally JG, et al. Genome-wide haploinsufficiency screen reveals a novel role for γ-TuSC in spindle organization and genome stability. Mol Biol Cell. 2013;24:2753–63.

52. Michaillat L, Mayer A. Identification of genes affecting vacuole membrane fragmentation in Saccharomyces cerevisiae. PLoS One. 2013;8:e54160.

53. Ivashov V, Zimmer J, Schwabl S, Kahlhofer J, Weys S, Gstir R, et al. Complementary α-arrestin-ubiquitin ligase complexes control nutrient transporter endocytosis in response to amino acids. Elife. 2020;9.

54. Dong K, Addinall SG, Lydall D, Rutherford JC. The yeast copper response is regulated by DNA damage. Mol Cell Biol. 2013;33:4041–50.

55. Burston HE, Maldonado-Báez L, Davey M, Montpetit B, Schluter C, Wendland B, et al. Regulators of yeast endocytosis identified by systematic quantitative analysis. J Cell Biol. 2009;185:1097–110.

56. Takuma T, Ushimaru T. Vacuolar fragmentation promotes fluxes of microautophagy and micronucleophagy but not of macroautophagy. Biochem Biophys Res Commun. 2022;614:161– 8.

57. Aitchison JD, Blobel G, Rout MP. Kap104p: a karyopherin involved in the nuclear transport of messenger RNA binding proteins. Science. 1996;274:624–7.

58. Lange A, Mills RE, Devine SE, Corbett AH. A PY-NLS nuclear targeting signal is required for nuclear localization and function of the Saccharomyces cerevisiae mRNA-binding protein Hrp1. J Biol Chem. 2008;283:12926–34.

59. Poirey R, Despons L, Leh V, Lafuente M-J, Potier S, Souciet J-L, et al. Functional analysis of the Saccharomyces cerevisiae DUP240 multigene family reveals membrane-associated proteins that are not essential for cell viability. Microbiology. 2002;148 Pt 7:2111–23.

60. Mukai H, Kuno T, Tanaka H, Hirata D, Miyakawa T, Tanaka C. Isolation and characterization of SSE1 and SSE2, new members of the yeast HSP70 multigene family. Gene. 1993;132:57–66.

61. Ho B, Baryshnikova A, Brown GW. Unification of Protein Abundance Datasets Yields a Quantitative Saccharomyces cerevisiae Proteome. Cell Syst. 2018;6:192–205.e3.

62. Hirata E, Shirai K, Kawaoka T, Sato K, Kodama F, Suzuki K. Atg15 in Saccharomyces cerevisiae consists of two functionally distinct domains. Mol Biol Cell. 2021;32:645–63.

63. Parzych KR, Ariosa A, Mari M, Klionsky DJ. A newly characterized vacuolar serine carboxypeptidase, Atg42/Ybr139w, is required for normal vacuole function and the terminal steps of autophagy in the yeast Saccharomyces cerevisiae. Mol Biol Cell. 2018;29:1089–99.

64. Van Den Hazel HB, Kielland-Brandt MC, Winther JR. Review: biosynthesis and function of yeast vacuolar proteases. Yeast. 1996;12:1–16.

65. Teter SA, Eggerton KP, Scott S V, Kim J, Fischer AM, Klionsky DJ. Degradation of lipid vesicles in the yeast vacuole requires function of Cvt17, a putative lipase. J Biol Chem. 2001;276:2083–7.

66. Spormann DO, Heim J, Wolf DH. Carboxypeptidase yscS: gene structure and function of the vacuolar enzyme. Eur J Biochem. 1991;197:399–405.

67. Wolf DH, Ehmann C. Studies on a proteinase B mutant of yeast. Eur J Biochem. 1979;98:375–84.

68. Longtine MS, McKenzie A, Demarini DJ, Shah NG, Wach A, Brachat A, et al. Additional modules for versatile and economical PCR-based gene deletion and modification in Saccharomyces cerevisiae. Yeast. 1998;14:953–61.

69. Sikorski RS, Hieter P. A system of shuttle vectors and yeast host strains designed for efficient manipulation of DNA in Saccharomyces cerevisiae. Genetics. 1989;122:19–27.

70. Gietz RD, Schiestl RH. Microtiter plate transformation using the LiAc/SS carrier DNA/PEG method. Nat Protoc. 2007;2:5–8.

71. Gietz RD, Schiestl RH. High-efficiency yeast transformation using the LiAc/SS carrier DNA/PEG method. Nat Protoc. 2007;2:31–4.

72. Alcalde M, Bulter T. Colorimetric assays for screening laccases. Methods Mol Biol. 2003;230:193–201.

73. Childs RE, Bardsley WG. The steady-state kinetics of peroxidase with 2,2’-azino-di-(3-ethyl-benzthiazoline-6-sulphonic acid) as chromogen. Biochem J. 1975;145:93–103.

74. Letunic I, Bork P. Interactive Tree of Life (iTOL) v6: recent updates to the phylogenetic tree display and annotation tool. Nucleic Acids Res. 2024;52:W78–82.

75. Duan S-F, Han P-J, Wang Q-M, Liu W-Q, Shi J-Y, Li K, et al. The origin and adaptive evolution of domesticated populations of yeast from Far East Asia. Nat Commun. 2018;9:2690.

76. Wang Q-M, Liu W-Q, Liti G, Wang S-A, Bai F-Y. Surprisingly diverged populations of Saccharomyces cerevisiae in natural environments remote from human activity. Mol Ecol. 2012;21:5404–17.

77. Fleige S, Pfaffl MW. RNA integrity and the effect on the real-time qRT-PCR performance. Mol Aspects Med. 2006;27:126–39.

78. Pazzagli M, Malentacchi F, Simi L, Orlando C, Wyrich R, Günther K, et al. SPIDIA-RNA: first external quality assessment for the pre-analytical phase of blood samples used for RNA based analyses. Methods. 2013;59:20–31.

79. Schmittgen TD, Livak KJ. Analyzing real-time PCR data by the comparative C(T) method. Nat Protoc. 2008;3:1101–8.

80. Widmer C, Lippert C, Weissbrod O, Fusi N, Kadie C, Davidson R, et al. Further improvements to linear mixed models for genome-wide association studies. Sci Rep. 2014;4:6874.

81. Ge SX, Jung D, Yao R. ShinyGO: a graphical gene-set enrichment tool for animals and plants. Bioinformatics. 2020;36:2628–9.

82. Xu S, Chen M, Feng T, Zhan L, Zhou L, Yu G. Use ggbreak to Effectively Utilize Plotting Space to Deal With Large Datasets and Outliers. Front Genet. 2021;12:774846.

83. Hughes CS, Moggridge S, Müller T, Sorensen PH, Morin GB, Krijgsveld J. Single-pot, solid-phase-enhanced sample preparation for proteomics experiments. Nat Protoc. 2019;14:68–85.

